# A PARP inhibitor, rucaparib, improves cardiac dysfunction in *ADP-ribose-acceptor hydrolase 3* (*Arh3*) deficiency

**DOI:** 10.1101/2023.02.07.527369

**Authors:** Sachiko Yamashita, Xiangning Bu, Hiroko Ishiwata-Endo, Jiro Kato, Danielle Springer, Audrey Noguchi, Morteza Peiravi, Chengyu Liu, Fan Zhang, Zu-Xi Yu, Randy Clevenger, Karen Keeran, Hong San, Martin J. Lizak, Joel Moss

## Abstract

**Aims:** Patients with *ADP-ribose-acceptor hydrolase 3* (*ARH3*) deficiency exhibit stress-induced childhood-onset neurodegeneration with ataxia and seizures (CONDSIAS). ARH3 degrades protein-linked poly(ADP-ribose) (PAR) synthesized by poly(ADP-ribose)polymerase (PARP)-1 during oxidative stress, leading to cleavage of the ADP-ribose linked to protein. *ARH3* deficiency leads to excess accumulation of PAR, resulting in PAR-dependent cell death or parthanatos. Approximately one-third of patients with homozygous mutant *ARH3* die from cardiac arrest, which has been described as neurogenic, suggesting that ARH3 may play an important role in maintaining myocardial function. To address this question, cardiac function was monitored in *Arh3*-knockout (KO) and - heterozygous (HT) mice.

**Methods and results:** *Arh3*-KO male mice displayed cardiac hypertrophy by histopathology and decreased cardiac contractility assessed by MRI. In addition, both genders of *Arh3*-KO and -HT mice showed decreased cardiac contractility by dobutamine stress test assessed by echocardiography. A direct role of ARH3 on myocardial function was seen with a Langendorff-perfused isolated heart model*. Arh3*-KO male mouse hearts showed decreased post-ischemic rate pressure products, increased size of ischemia-reperfusion (IR) infarcts, and elevated PAR levels. Consistently, *in vivo* IR injury showed enhanced infarct size in *Arh3*-KO mice in both genders. In addition, *Arh3*-HT male mice showed increased size of *in vivo* IR infarcts. Treatment with an FDA-approved PARP inhibitor, rucaparib, improved cardiac contractility during dobutamine-induced stress and exhibited reduced size of *in vivo* IR infarcts. To understand better the role of ARH3, CRISPR-Cas9 was used to generate different *Arh3* genotypes of myoblasts and myotubes. Incubation with H2O2 decreased viability of *Arh3*-KO and -HT myoblasts and myotubes, resulting in PAR-dependent cell death that was reduced by PARP inhibitors or by transfection with the *Arh3* gene.

**Conclusion:** ARH3 regulates PAR homeostasis in myocardium to preserve function and protect against oxidative stress; PARP inhibitors reduce the myocardial dysfunction seen with *Arh3* mutations.

**Graphical Abstract:** 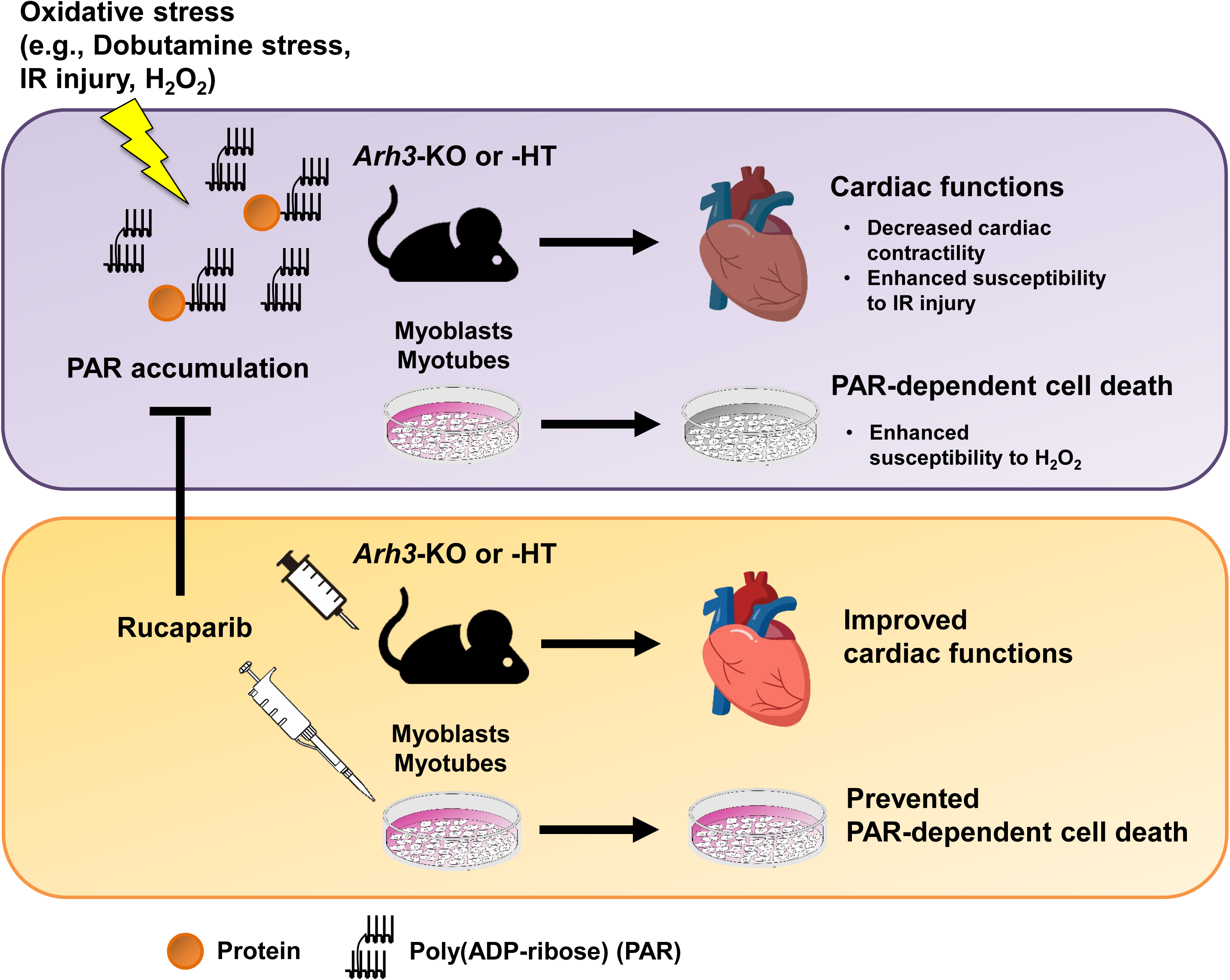

## 1. Introduction

Poly-ADP-ribosylation (PARylation) is a reversible post-translational modification, which is involved in DNA repair, transcription control, and cell death.^1, 2^ In PARylation, poly(ADP-ribose) polymerases (PARPs) transfer the ADP-ribose moiety of NAD^+^ to protein, forming a branched, long-chain polymer.^1, 2^ In response to oxidative stress, nuclear PARP-1 catalyzes the synthesis of poly(ADP-ribose) polymers (PARs), which are covalently linked to specific amino acid residues (e.g., glutamate, aspartate, lysine, arginine, serine) of acceptor proteins, including automodification of PARP-1 itself.^1, 2^ Excessive activation of PARP-1 caused by oxidative stress is seen in several cardiovascular diseases (e.g. atherosclerosis, myocardial hypertrophy, cardiomyopathy, coronary artery disease, hypertension, several forms of heart failure), with the underlying mechanism still unclear.^3–5^

ADP-ribose-acceptor hydrolase 3 (ARH3), a member of the 39-kDa ARH family (ARH1-3), hydrolyzes PAR in an exo-glycosidic reaction, generating free ADP-ribose and PAR oligomers in a Mg^2+^-dependent cleavage reaction.^6–10^ In addition, ARH3 also hydrolyzes ADP-ribosyl- (serine) protein α-NAD^+^ and *O*-acetyl-ADP-ribose (*O*AADPr), which is synthesized by Sir2 family of enzymes by NAD^+^-dependent deacetylation of proteins.^7, 8^ ARH3 is located in nuclei, cytoplasm, and mitochondria.^9^ PAR glycohydrolase (PARG) is the primary protein hydrolyzing PAR, catalyzing the cleavage of PAR chains to yield ADP-ribose and PAR oligomers through exo-glycosidic and endo-glycosidic reactions.^10^ Different mechanisms of PAR recognition and the cellular localization of PAR and ARH3 appear to be responsible for the unique cellular roles of ARH3 and PARG.

*ARH3* deficiency results in a rare, autosomal-recessive, multi-organ syndrome, which is characterized by stress-induced childhood-onset neurodegeneration with ataxia and seizures (CONDSIAS).^11–17^ Patients may show developmental delays or intellectual impairment, gait abnormalities, neuropathy, facial myoclonia, sensorineural hearing loss, and ophthalmologic disorders.^11–17^ Using fibroblasts from patients and mice with ARH3 deficiency, we previously showed that ARH3 deficiency increased PAR accumulation in the nucleus and cytoplasm following oxidative stress, inducing translocation of apoptosis-inducing factor (AIF) from mitochondria to the nucleus, triggering chromatin condensation, activation of endo-nucleases and large-scale DNA fragmentation, resulting in PAR (or PARP-1)-mediated cell death or parthanatos.^17, 18^ This cell death pathway is induced in patients with *ARH3* deficiency and is the pathogenesis of CONDSIAS.^11–17^ Recently, a patient with heterozygous (HT) mutation in ARH3 showed autonomic nervous dysfunction such as polyuria, gastrointestinal disturbance, sinus arrhythmia, and myogenic lesions as additional phenotypic features.^16^ In addition, about one-third of patients with *ARH3* deficiency died from neurogenic cardiac arrest.^11, 13–15^ We hypothesized that these cardiac abnormalities resulted from an intrinsic myocardial *ARH3* mutation. To test this hypothesis, we generated *Arh3*-knockout (KO), - HT, and wild-type (WT) mice as well as the corresponding C2C12 myoblasts and myotubes to determine the effects of *ARH3* deficiency on the murine heart and cultured murine skeletal muscle cells.

In this report, we describe, in *Arh3*-KO mice, cardiac hypertrophy, reduced the percentage of ejection fraction (%EF) using MRI, reduced %EF and fractional shortening during dobutamine infusion using echocardiography, and enhanced *in vivo* and *ex vivo* (Langendorff) myocardial ischemia-reperfusion (IR) injury. Similar to *Arh3*-KO mice, *Arh3*-HT mice showed reduced cardiac contractility during dobutamine infusion and increased *in vivo* IR infarcts. We hypothesized that prevention of increased PAR levels by PARP inhibition would improve the cardiac dysfunction and IR injury. In prior reports of effects of PARP inhibitors on myocardial function, classical PARP inhibitors (e.g., 3- aminobenzamide, PJ34, INO-1001) improved cardiac contractility and enhanced recovery from myocardial IR injury in mouse, rat, rabbit, dog, and pig models.^3, 19–21^ We observed that an FDA-approved PARP inhibitor, rucaparib, improved cardiac contractility during dobutamine stress and reduced the size of *in vivo* IR infarcts in *Arh3*-KO mice. At the cellular level, incubation with H2O2 decreased viability of *Arh3*-KO and -HT myoblasts/myotubes compared to WT, resulting in PAR-dependent cell death that was reduced by PARP inhibitors or by transfection with the *Arh3* gene. These results indicate that *Arh3* regulates myocardial PAR levels to protect cardiac function under oxidative stress, preventing PAR-dependent cell death.

## 2. Methods

### 2.1 Generation of *Arh3*-KO and -HT mice

Generation of *Arh3-*KO and -HT mice was described by Mashimo et al.^17^ *Arh3-*KO and -HT mice were backcrossed for 9 generation to the C57BL6/L background.^17^ Genotypes of these mice were confirmed by PCR.^17^

### 2.2 Histological sections

Mouse hearts were fixed with formalin and embedded in paraffin. Myocardium sections were stained with hematoxylin and eosin. Representative images of 13 months old *Arh3*-WT, -HT, and -KO mouse hearts are shown in *Figure 1B and C*.

**Figure 1.**
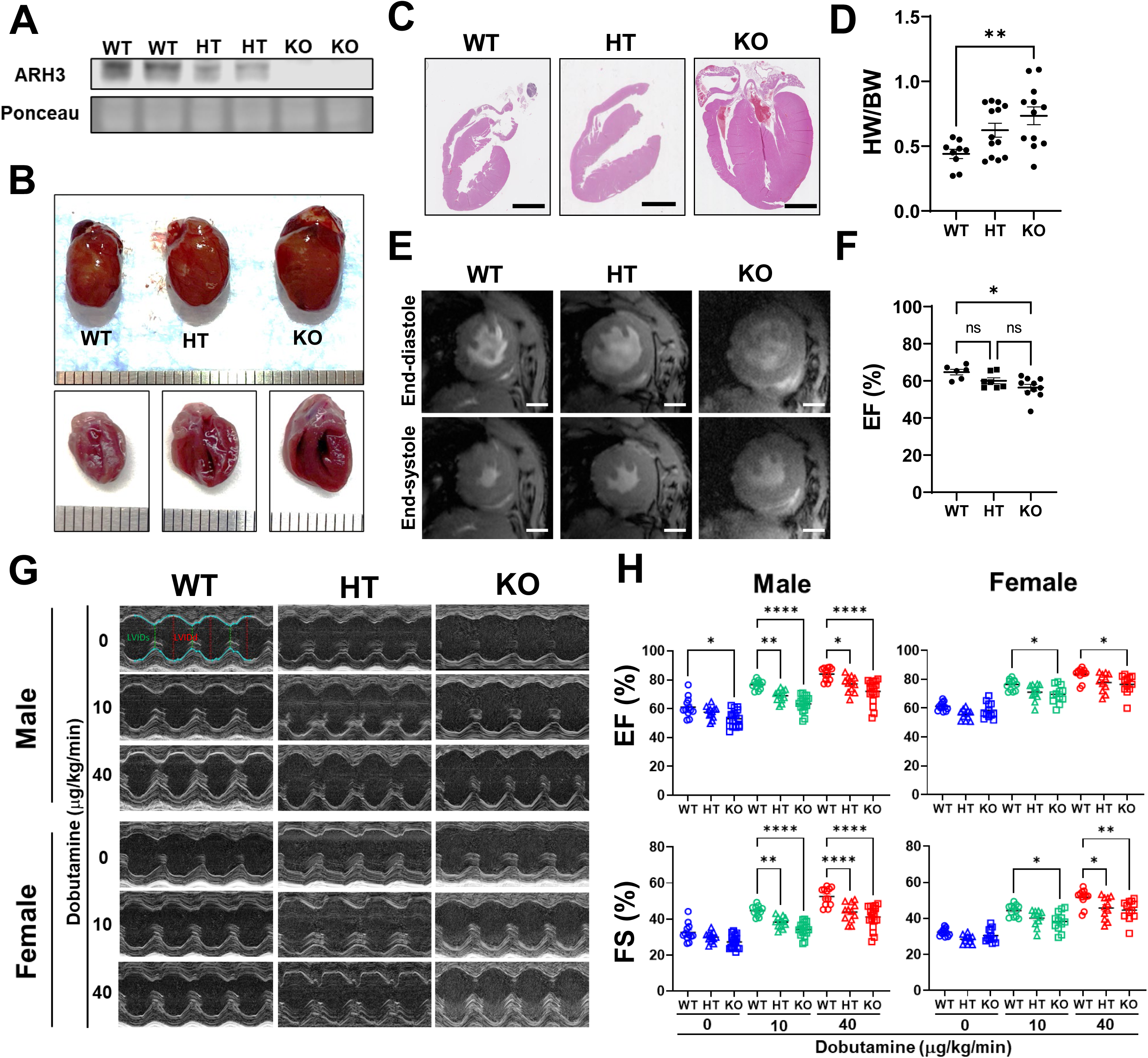
*Arh3*-KO and -HT mice showed reduced cardiac contractility in both genders under dobutamine stress. (*A*) Endogenous ARH3 protein levels in mouse heart among genotypes. Tissues were subjected to Western blotting using anti-ARH3 antibody. (*B*) Representative of gross heart images of wild-type (WT), *Arh3*-heterozygous (HT), and *Arh3*- knockout (KO) mice. Age: 13-month-old. (*C*) Hematoxylin and eosin staining of WT, *Arh3*- HT, and *Arh3*-KO mouse heart. *Arh3*-KO mouse heart shows hypertrophy. (Scale bar, 2 mm), Age: 13-month-old. (*D*) The heart weight per body weight ratio among genotypes. Two-way ANOVA Tukey’s multiple comparisons test, \*\**p* < 0.01, Age: WT males, *n* = 9, 47.5 ± 3.3 weeks; HT males, *n* = 13, 47.1 ± 3.4 weeks; KO males, *n* = 12, 49.2 ± 1.5 weeks. (*E*) Representative MRI images among *Arh3* mouse genotypes. (Scale bar, 2 mm) (*F*) Ejection fraction by MRI in male mice of different *Arh3* genotypes. Two-way ANOVA Tukey’s multiple comparisons test, \**p* < 0.05, Age: WT males, *n* = 6, 40.8 ± 3.9 weeks; HT males, *n* = 7, 42.4 ± 5.4 weeks; KO males, *n* = 10, 49.2 ± 9.1 weeks. (*G*) Representative echocardiographic images among *Arh3* mouse genotypes subjected to dobutamine stress. (*H*) Assessments of %ejection fraction (EF) and fractional shortening (FS) in male and female mice of different *Arh3* genotypes, subjected to dobutamine stress (0, 10, and 40 μg/kg/min). Two-way ANOVA Tukey’s multiple comparisons test, \**p* < 0.05, \*\**p* < 0.01, \*\*\**p* < 0.001, \*\*\*\**p* < 0.0001. Dobutamine stress test was performed in WT males, *n* = 12, 37.2 ± 4.1 weeks; HT males, *n* = 12, 37.1 ± 2.1 weeks; KO males, *n* = 18, 39.1 ± 3.4 weeks; WT females, *n* = 12, 36.1 ± 2.3 weeks; HT males, *n* = 11, 35.9 ± 1.7 weeks; KO females, *n* = 12, 37.4 ± 2.9 weeks.

### 2.3 Magnetic resonance imaging (MRI)

The mouse was anesthetized with 2-3% isoflurane and maintained with 1.0-2.5% isoflurane in a medical air/oxygen mixture. The mouse was placed prone on a plastic cradle. The temperature, breath rate, and heart rate were monitored throughout the experiment for maintenance of anesthesia. MRI imaging was performed on a 9.4 Tesla Bruker Neo HD MRI scanner. Following acquisition of anatomical locator images, a series of 1mm slices was prescribed axially, in sequence, along the heart chambers. Cine imaging was performed using the Bruker retrospectively gated cardiac sequence (INTRAGATE). The parameters for imaging were FOV = 25 mm x 25 mm, matrix = 256 x 256, slice thickness=1mm, TR/TE = 91/3.7 ms. Sixteen movie frames were acquired and oversampling was set so that image acquisition time was approximately 30 minutes for 5 slices. The images were reconstructed using the Bruker proprietary software and then exported as DICOM for analysis.

### 2.4 Dobutamine stress with echocardiography

Dobutamine stress testing was performed using 10-month-old mice. Mice were placed on a heated imaging platform with electrocardiography leads and lightly anesthetized with 2% isoflurane via a nose cone during echocardiography and dobutamine infusions. Dobutamine was administered by intravenous infusion via a tail-vein catheter. After the baseline-scan, the mice received constant rate infusions of dobutamine (0.625 mg/mL in normal saline containing 5% dextrose). The first administration, a low dose, was 10 mg/kg/min. After the heart rate reached a steady state approximately 500 beats/min, a higher dose (40 mg/kg/min) was given. The length of time of the infusions was between approximately 20 and 30 min. Cardiac images were obtained using the Vevo2100 ultrasound system (FUJIFILM VisualSonics Inc., Toronto, Canada) with a 30-MHz ultrasound probe at baseline and during the low and high dose dobutamine infusions. For the treatment assay, PARP inhibitor rucaparib (MedChemExpress) at 10 mg/kg or vehicle buffer (1.25% Dimethyl sulfoxide (DMSO), 10% 2- hydroxypropyl-b-cyclodextrin (Sigma-Aldrich), 0.4% sodium chloride) was administered by intraperitoneal injection two times, 24 h and 1 h before dobutamine stress.

### 2.5 A Langendorff-perfused isolated heart model of ischemia-reperfusion injury

Ten-month old mice were subjected to a Langendorff heart model of ischemia reperfusion. Mice were anesthetized with 50 mg/kg pentobarbital via intraperitoneal injection and then anticoagulated with 50 μL of heparin (1000 U/ml) via intraperitoneal injection. Hearts were harvested and immediately placed in ice-cold Krebs-Henseleit (KH) buffer (120 mM NaCl, 11 mM D-glucose, 25 mM NaHCO3, 1.75 mM CaCl2, 4.7 mM KCl, 1.2 mM MgSO4, and 1.2 mM KH2PO4). Hearts were cannulated and suspended in the Langendorff apparatus. Then, the hearts were retrogradely perfused with oxygenated solution KH buffer (95% O2 and 5% CO2) to maintain pH at 7.4 at a constant pressure of 100 cm H2O at 37°C. As shown in *Figure 3A-D*, isolated hearts were subjected to 5 min of perfusion, and then 25 min of ischemia before 30 or 90 min of reperfusion. Control hearts were immediately harvested after 5 min of perfusion.

### 2.6 *In vivo* myocardial ischemia-reperfusion injury

Ten-month old mice were subjected to acute myocardial infarction (MI) surgery by surgeons who were blinded to the mouse genotype. Mice were anesthetized with 1-3% isoflurane and positioned in dorsal recumbency. The mice were intubated by making a midline skin incision over the neck, bluntly separating the sternohyoid muscle to view the trachea, and passing a 20g angiocatheter through the mouth into the trachea. After intubation, the mouse was tilted onto its right side and the skin incision was extended midline to just below the xiphoid process. The left thorax muscles were retracted to expose the left ribcage. The chest was entered at the 3^rd^ intercostal space (between the 3^rd^ and 4^th^ ribs). Micro-scissors were used to cut the intercostal muscle along the cranial aspect of the 4^th^ rib. The ribs were retracted to view the left side of the heart. The pericardium was opened. At approximately one-third the length of heart from the apex, a 7-0 suture was placed around the left anterior descending artery (LAD) and tied with a slipknot to occlude the vessel. A darkening of the lower left ventricle extending to the apex was confirmation of occlusion. To ensure the LAD remained occluded, the chest was kept open for periodically checking the color of the infarct area. Warm saline-moistened gauze was placed over the incision to keep the tissues hydrated. The slipknot was released when reperfusion was achieved and a lightening color change was used to indicate reperfusion. The chest was closed after reperfusion was observed. For the treatment assay, the PARP inhibitor, rucaparib, at 10 mg/kg or control buffer (2.5% dimethyl sulfoxide, 10% 2-hydroxypropyl-b-cyclodextrin, 0.4% sodium chloride) was administered by intraperitoneal injection 4 times at indicated time points: 1 day before ischemia, 1 h before ischemia, 10 min into ischemia, and 30 min into reperfusion. After 24-h reperfusion, mice were anesthetized, and 1 ml of 1% potassium chloride solution was administered via abdominal inferior vena cava to cause cardiac arrest. The heart was excised and sectioned in 2-mm slices and then stained with 1% triphenyl tetrazolium chloride (TTC) (Millipore Sigma) in PBS solution at 37°C for 20 min. These heart slices were fixed in 10% buffered formalin to quantify the infarct size by ImageJ.

### 2.7 Reagents and antibodies

Rucaparib and olaparib were purchased from MedChemExpress; mouse monoclonal anti-poly(ADP-ribose) (PAR) (10H) antibodies from Enzo Life Sciences; mouse monoclonal anti-DDK antibodies from Origene; mouse monoclonal anti-myogenin antibodies from BD Biosciences; rabbit polyclonal anti-mouse ADP-ribosyl-acceptor hydrolase (ARH3) antibodies were produced in rabbit by injecting ARH3 peptide (YenZym antibodies LLC). Additional details are provided in online Supplementary material.

### 2.8 Cells

C2C12 cells and MEFs were cultured in Dulbecco’s Modified Eagle Medium (DMEM) supplemented with 10% heat-inactivated fetal bovine serum (FBS), penicillin (100 U/ml) and streptomycin (100 μg/ml) in a humidified incubator with 5% CO2 at 37.0°C. Detailed protocols are described in online Supplementary material.

### 2.9 Using CRISPR/Cas9 to generate C2C12-*Arh3* null and haploinsufficient mutants

The *Arh3* KO and HT mutant C2C12 cell lines were generated using CRISPR/Cas9. Detailed protocols are presented in online Supplementary material.

### 2.10 Cytotoxicity assays

C2C12 myoblasts and MEFs (1.0 × 10^4^ or 3 × 10^3^ cells) were incubated on 96-well plates for 1 day before exposure for 3 h to the indicated concentrations (0.1-1 mM) of H2O2. C2C12 myotubes (1.2 × 10^4^ cells) were incubated on 96-well plates for 1 day and then, media were changed to 2% horse serum and culture was continued for 5 days before exposure for 3 h to the indicated concentrations (0.1-1 mM) of H2O2. PARP inhibitors rucaparib (0.1-10 μM) and olaparib (0.1-10 μM) were added for 30 min prior to H2O2 exposure. Cell viability was assessed using a Cell Counting Kit-8 (Dojindo). Detailed methods are available in online Supplementary material.

### 2.11 SDS-PAGE and Western blotting

C2C12 myoblasts and MEFs (5.0 × 10^5^ cells) were incubated on 6-well plates for 1 day before exposure for 3 h to 0.3 mM of H2O2. Cells were lysed in 20 mM Tris-HCl (pH 7.4) with 2% SDS supplemented with a proteinase inhibitor (cOmplete^TM^, EDTA-free Protease Inhibitor Cocktail, Roche). Protein concentration of lysates was measured using a bicinchoninic acid (BCA) assay kit (Thermo Scientific). Each sample was subjected to Bis-Tris SDS-PAGE (Invitrogen) and then transferred to nitrocellulose membranes (Invitrogen). Each membrane was blocked with Odyssey Blocking Buffer (LI-COR) or 2 % (w/v) Blocking-Grade Blocker (BIO-RAD) at room temp for 1 h and then incubated with primary antibody at 4°C overnight. Each membrane was washed with 1 x Tris-buffered saline containing 0.05% Tween 20 and then incubated with an appropriate secondary antibody (LI-COR) in blocking buffer for 1 h at room temperature. Each membrane was washed again with Tris-buffered saline and analyzed by an Odyssey Imaging Systems (LI-COR).

### 2.12 Data analysis and statistics

Statistical analysis was performed using a GraphPad Prism or Microsoft Excel software. Statistical comparisons analysis between 2 groups were assessed by paired *t* test. Statistical Multiple comparisons of more than 2 groups were assessed by two-way ANOVA Bonferroni’s multiple comparison test or two-way ANOVA Tukey’s multiple comparisons test. *P* values less than 0.05 were significant.

### 2.13 Study approval

Experimental protocols for mouse studies (H-0128R5, H-0271R3, and H-0172R5) were approved by the National Heart, Lung, and Blood Institute (NHLBI) Animal Care and Use Committee. Studies involving the *ARH3*-deficient family were approved under NHLBI Institutional Review Board (IRB) protocol 96-H-0100.

## 3. Results

### 3.1 *Arh3*-KO and -HT mice showed reduced motor function and coordination in an age-dependent manner assessed by open-field and rotarod tests

Previously, presumed cardiac abnormalities seen in patients with *ARH3* deficiency were believed to result from neurological dysfunction.^11–17^ To investigate whether *Arh3* deficiency alters motor function, we assessed locomotor activity, motor coordination, and muscle strength using the open-field, rotarod, and isometric-torque tests, respectively. In the open-field test, assessing voluntary movement among 3-4 months old mice, no difference was observed in total distance traveled in 30 min by mice, regardless of gender and genotype (*Figure S1A*, left). However, in male 7-8-month old mice, the total distance traveled by *Arh3*-KO and *Arh3*-HT mice was significantly less than that of WT mice (*Figure S1A*, right). In addition, in female 7-8-month old mice, the total distance traveled by *Arh3*-KO mice was significantly less than that of WT and HT mice (*Figure S1A*, right). Thus, older mice with *Arh3* mutations exhibited reduced voluntary lower locomotor activity (*Figure S1A*). Extended run time in the rotarod test shows coordinated running, consistent with normal neurological activity. The total run time in 3-4 months old *Arh3*-KO and -HT mice was significantly reduced from that of WT mice regardless of gender (*Figure S1B*, left). Although the total run time in 7-8 months old *Arh3*-KO male mice was significantly reduced from that of WT, there was no significant difference between male WT and HT mice (*Figure S1B*, right). In addition, no significant difference among genotypes in female mice was observed (*Figure S1B*, right). Summary table of results are shown in *table S1*. Since isometric torque test involving skeletal muscle of hind legs of *Arh3*-KO and -HT mice was not significantly different from WT mice regardless of gender (*Figure S1C*), the results of the open-field test in 7-8 months old mice and the rotarod test in 3-4 months old *Arh3*-KO and -HT mice might be due to neurological dysfunction resulting from the *Arh3* mutation. However, the results of the rotarod test in 7-8 months old mice could not be explained primarily by neurological abnormalities, and cardiac dysfunction might be responsible for some of the decreased locomotor activity. To analyze the direct function of *Arh3* mutation on the heart, the effect of *Arh3* mutation on the structure and physiological function of the heart were analyzed as described below.

### 3.2 *Arh3*-KO and -HT mice displayed decreased cardiac contractility regardless of gender assessed by MRI and echocardiography

To study effects of *Arh3* mutations related to cardiac dysfunction, we used *Arh3*-KO, -HT, and -WT mice. According to the Human Protein Atlas, ARH3 protein is moderately expressed in human cardiomyocytes (https://www.proteinatlas.org/ENSG00000116863-ADPRHL2). ARH3 protein was identified in the WT mouse heart using Western blotting analysis (*Figure 1A*). As expected, ARH3 protein was expressed at lower levels in the HT mouse heart compared to the WT heart (*Figure 1A*). Expression was not observed in the KO male mouse heart (*Figure 1A*). Of interest, representative gross heart images and histological sections indicated hypertrophy in the *Arh3*-KO male heart (3 hypertrophies/10 hearts), compared to the *Arh3*-HT (0 hypertrophy/11 hearts) and WT mouse male hearts (0 hypertrophy/10 hearts) (*Figure 1B and C*). These findings in murine *Arh3-*KO hearts are consistent with observations in a patient with *ARH3* deficiency who had left ventricular hypertrophy and mitral insufficiency.^15^ In addition, the heart weight/body weight (HW/BW) ratio in the *Arh3*-KO mice was significantly higher than that of the WT mice; moreover, the *Arh3*-HT mice trended to have a higher HW/BW ratio than the WT mice (*p* = 0.086) (*Figure 1D*). To investigate the role of *Arh3* in cardiac function, we assessed cardiac contractility in mice by MRI or with a dobutamine stress test by echocardiography. We monitored cardiac contractility in 10-months-old mice by MRI. Representative images of short-axis in the endo-diastole and -systole, and video of cardiac contractility among genotypes are shown in *Figure 1E* and online Supplementary material, *Video S1*. The percentage of ejection fraction (%EF) was significantly less in the *Arh3*-KO mice than the WT male mice (*Figure 1F*). Thus, *Arh3*-KO mice showed decreased cardiac contractility compared to the WT at baseline by MRI. In a dobutamine stress test by echocardiography, we administered dobutamine (0, 10, and 40 μg/kg/min) to mice, age ∼ 10 months. Representative images of echocardiography under dobutamine infusion are shown in *Figure 1G*. With dobutamine infusion, cardiac contractility increased regardless of genotype or gender in a dose- dependent manner (*Figure 1G and H, Figure S2*). In the male mice, the %EF was significantly less in the *Arh3*-KO mice than in the WT mice at baseline (*Figure 1H*, upper left). The left ventricular internal dimension (LVID) at diastole (LVIDd) and systole (LVIDs) in the *Arh3*- KO mice were significantly larger than that in the WT mice at baseline (*Figure S2*, left). The %EF and fractional shortening (FS) were significantly less in the *Arh3*-KO mice than in the WT mice, whether they received a low-dose dobutamine infusion (10 μg/kg/min) or high-dose dobutamine infusion (40 μg/kg/min) (*Figure 1H*, left). In addition, %EF and %FS in the *Arh3*-HT mice were significantly less than in the WT mice with either the low- or high-dose dobutamine infusion (*Figure 1H*, left). The LVIDd and LVIDs in the *Arh3*-KO mice were significantly larger than in the WT mice, with either low- or high-dose dobutamine infusion (*Figure S2*, left). The LVIDs in the *Arh3*-HT mice was significantly larger than in the WT mice with low-dose dobutamine infusion (*Figure S2*, left). In the female mice, no significant difference in cardiac parameters was observed regardless of genotypes at baseline. The % EF and %FS in the *Arh3*-KO mice were significantly less than in the WT mice with either the low- or high-dose dobutamine infusion (*Figure 1H*, right). In addition, %FS in the *Arh3*-HT mice was significantly less than in the WT mice on the high-dose dobutamine infusion (*Figure 1H*, right). The LVIDs in *Arh3*-KO mice was significantly larger than in the WT mice with the high-dose dobutamine infusion (*Figure S2*, right). The stroke volume (SV) and heart rate (HR) were not significantly different regardless of gender or genotype (*Figure S2*, right). These results suggest that *Arh3*- KO and -HT mice displayed reduced cardiac contractility with dobutamine stress regardless of gender.

### 3.3 PARP inhibitor, rucaparib, improves decreased cardiac contractility in *Arh3*-KO and -HT mice

It has been reported that PARP inhibitors attenuate the suppression of cardiac contractility elicited by doxorubicin or endotoxin in murine models of heart failure.^20^ To investigate whether a PARP inhibitor improves the cardiac dysfunction seen with dobutamine infusion, we used the FDA-approved PARP inhibitor rucaparib in male mice. Ten-month-old mice were injected with control buffer (5% DMSO, 10% 2- hydroxypropyl-β-cyclodextrin, 0.4% sodium chloride) or 10 mg/kg of rucaparib via intraperitoneal injection 1 day and 1 h before a dobutamine stress test. At baseline, rucaparib increased %EF and %FS in *Arh3*-KO mice, resulting in no significant differences between WT and KO mice (*Figure 2A*). Rucaparib also significantly increased %EF and %FS, and decreased LVIDs in *Arh3*-KO mice, whether they received a low- or high-dose dobutamine infusion (*Figure 2A* and *Figure S3*, left). Similar to the results seen in *Arh3*-KO mice, rucaparib significantly increased %EF in *Arh3*-HT mice, whether they received a low- or high-dose dobutamine infusion (*Figure 2B*). Of interest, rucaparib increased %EF (Trend, *p* = 0.091) and significantly increased %FS (p < 0.05) in *Arh3*-WT mice with low-dose dobutamine infusion (*Figure 2*). Thus, PARP inhibitor, rucaparib improved decreased cardiac contractility in *Arh3*-KO, -HT mice, and -WT mice.

**Figure 2.**
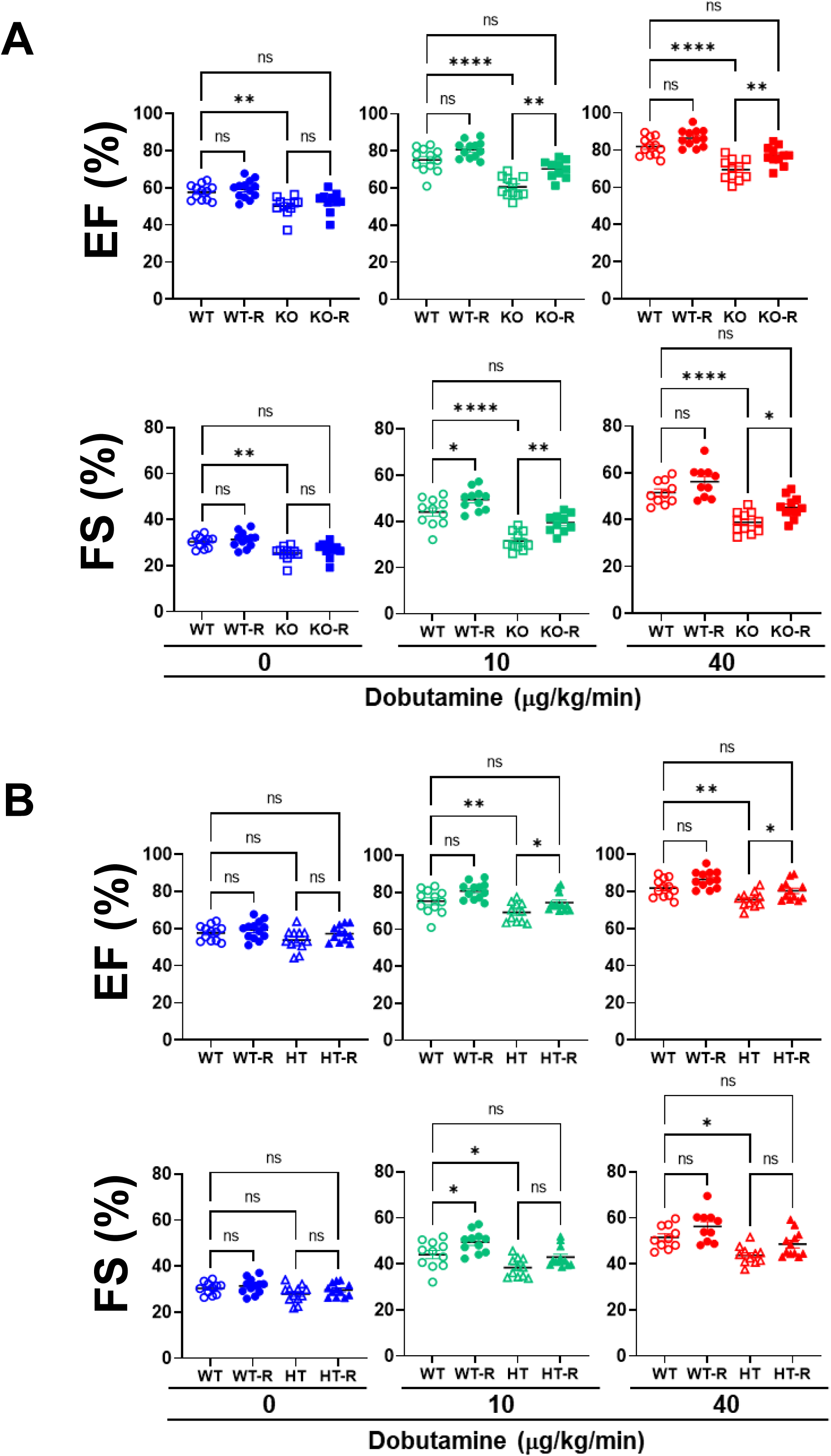
Rucaparib improved decreased cardiac contractility in *Arh3*-KO and -HT mice. (*A*) Assessments of % of EF and FS in *Arh3*-WT and -KO male mice subjected to dobutamine stress (0, 10, and 40 μg/kg/min) with or without 10 mg/kg rucaparib. Two-way ANOVA Tukey’s multiple comparisons test, ns: no significant difference, \**p* < 0.05, \*\**p* < 0.01, \*\*\*\**p* < 0.0001, Dobutamine stress test was performed in WT+DMSO males, *n* = 11, 41.0 ± 4.5 weeks; WT + rucaparib males, *n* = 11, 41.0 ± 5.2 weeks; KO + DMSO males, *n* = 11, 41.8 ± 6.0 weeks; KO + rucaparib males, *n* = 11, 39.3 ± 6.2 weeks. (*B*) Assessments of %EF and FS in *Arh3*-WT and -HT male mice subjected to dobutamine stress with or without 10 mg/kg rucaparib. Two-way ANOVA Tukey’s multiple comparisons test, ns: no significant difference, \**p* < 0.05, \*\**p* < 0.01. Dobutamine stress test was performed in WT + DMSO males, *n* = 11, 41.0 ± 4.5 weeks; WT + rucaparib males, *n* = 11, 41.0 ± 5.2 weeks; HT + DMSO males, *n* = 12, 39.5 ± 7.3 weeks; HT+ rucaparib males, *n* = 12, 41.1 ± 7.9 weeks.

### 3.4 *Arh3*-KO male mice showed increased myocardial ischemia-reperfusion injury in the *ex vivo* Langendorff-perfused isolated heart model

Clinical data show that patient families with *ARH3* deficiency have experienced repeated cardiac arrest and sudden death.^11, 13–15^ To determine the myocardial effect of *Arh3*-KO, we examined IR injury in 10-months old mice. To evaluate the direct effects of ARH3 on cardiac function without involvement of neurological and humoral factors, mouse hearts were examined using the *ex vivo* Langendorff-perfused isolated heart model. In this *ex vivo* model, the heart function recovery rate (post-ischemic rate pressure products) of the *Arh3-*KO mouse hearts (27.56% ± 1.38%, *n* = 8) was significantly lower than that of the WT mouse hearts (44.15 ± 1.53%, *n* = 8) with 25-min ischemia and 90- min of reperfusion (*Figure 3A*). In addition, the infarct size of *Arh3*- KO (61.91 ± 0.91%, *n* = 8) mice was significantly larger than that seen in the WT mice (39.13 ± 1.11%, *n* = 8) after reperfusion (*Figure 3B and C*). To investigate the hypothesis that PAR accumulation in *Arh3*-KO mouse hearts increases during IR injury, post-IR injury hearts were subjected to Western blotting using anti-PAR-antibody. Consistent with results of IR injury, PAR levels of the *Arh3*-KO mouse hearts were higher than in WT mice after 25-min ischemia and 30-min reperfusion (*Figure 3D*). Thus, ARH3 regulates PAR metabolism during IR injury in the Langendorff model.

**Figure 3.**
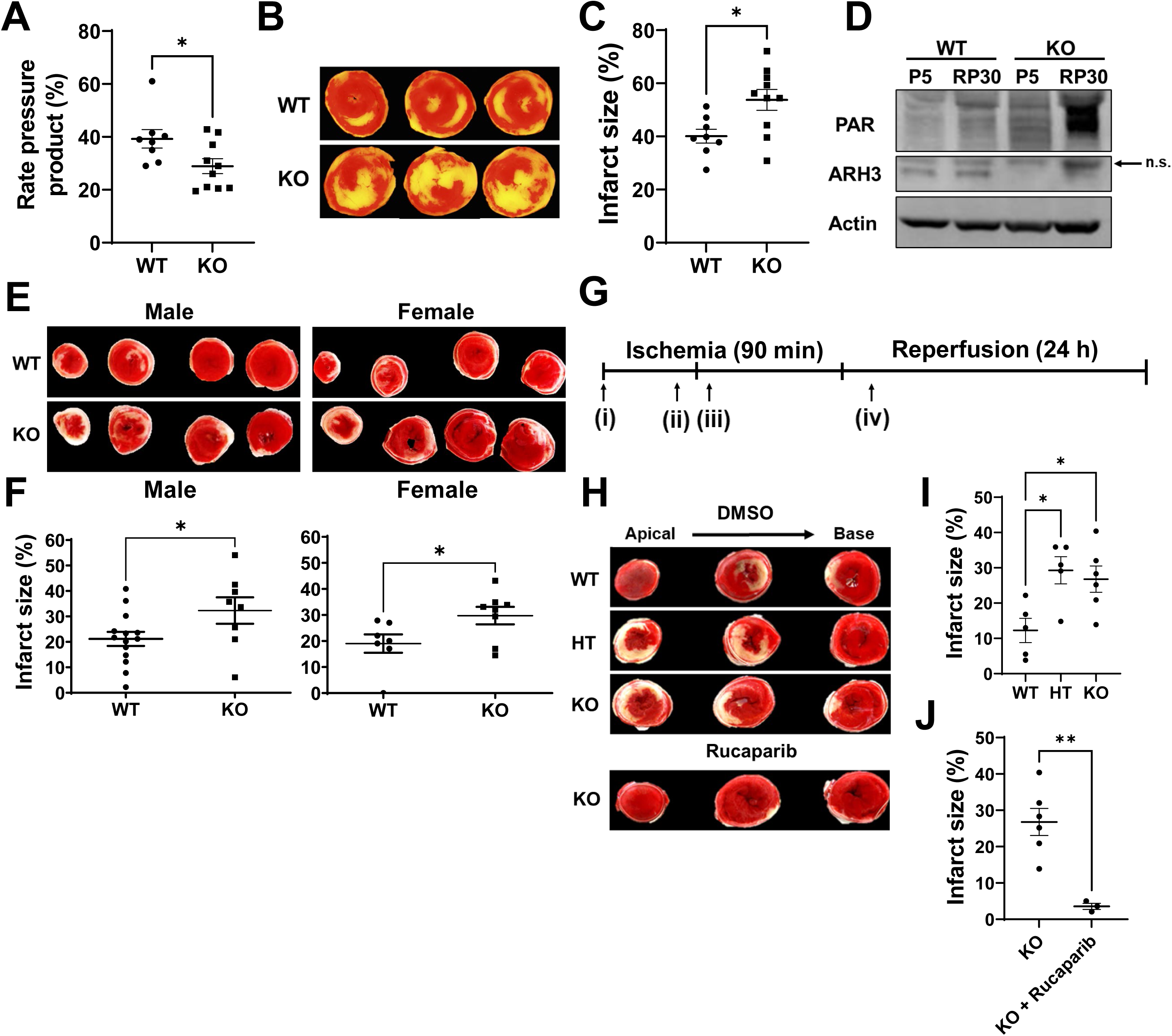
*Arh3*-KO and -HT mice showed increased ischemia-reperfusion- induced myocardial injury that is prevented by rucaparib. (*A*) Recovery of post-ischemic left ventricular (Rate pressure product) after *ex vivo* IR injury. T-test, \**p* < 0.05. E*x vivo* IR injury using a Langendorff-perfused isolated heart model was performed in WT males, *n* = 9, 39.9 ± 6.4 weeks; KO males, *n* = 10, 44.8 ± 9.3 weeks. (*B*) Representative pictures of male mouse hearts in *Arh3*-WT and -KO subjected to 20 minutes perfusion, 25 minutes ischemia, and 90 minutes reperfusion injury (Langendorff-perfused isolated heart model). Triphenyl tetrazolium chloride (TTC) staining shows the infarct area (Yellow) of mouse heart. (*C*) Quantification of the infarct size of (*B*). T-test, \**p* < 0.05. (*D*) PAR and ARH3 protein levels in male mouse hearts. Heart extracts were subjected to Western blotting using anti-poly(ADP-ribose) (10H) antibody or anti-ARH3 antibody or anti-actin antibody as an internal control. P5, perfusion 5 min; RP30, ischemia 25 min followed by reperfusion 30 min. An upper band reactive with anti-ARH3 antibody is nonspecific (n.s.). (*E*) Representative pictures of male and female mouse hearts in *Arh3*-WT and -KO subjected to 2-h ischemia and 24-h reperfusion injury (*in vivo* myocardial ischemia-reperfusion). TTC staining shows the infarct area (white) of mouse heart. (*F*) Quantification of the infarct size of (*E*). T-test, \**p* < 0.05. *In vivo* IR injury was performed in WT males, *n* = 14, 41.1 ± 4.27 weeks; KO males, *n* = 8, 39.4 ± 7.55 weeks; WT females, *n* = 7, 52.1 ± 4.34 weeks; KO males, *n* = 8, 52.0 ± 7.68 weeks. (*G*) The ischemia-reperfusion protocol. Mouse heart was subjected to 90 min ischemia and 24 h reperfusion with or without control (DMSO) or 10 mg/kg rucaparib. Injection time points are (i) 1 day before ischemia; (ii) 1 h before ischemia; (iii) 10 min into ischemia; (iv) 30 min into reperfusion. (*H*) Representative pictures of male mouse heart between genotypes treated with control (DMSO) or 10 mg/kg rucaparib during ischemia-reperfusion injury. TTC staining shows the infarct area (white) of mouse heart. (I) Quantification of the infarct size of *(H)*. The infarct size in *Arh3*-HT or -KO male mouse heart was significantly larger than that in the WT male mice. Two-way ANOVA Bonferroni’s multiple comparisons test, \**p* < 0.05, *In vivo* IR injury was performed in WT males, *n* = 5, 41.4 ± 1.1 weeks; HT males, *n* = 5, 41.6 ± 1.5 weeks; KO males, *n* = 6, 40.6 ± 4.0 weeks. (*J*) Quantification of the infarct size of *(H)* treated with rucaparib. Ten mg/kg rucaparib significantly reduced the infarct size of male *Arh3*-KO mouse heart compared to control. Two-way ANOVA Bonferroni’s multiple comparisons test, \*\**p* < 0.01, *In vivo* IR injury was performed in KO + DMSO males, *n* = 6, 40.6 ± 4.0 weeks; KO + rucaparib males, *n* = 3, 39.2 ± 0.4 weeks.

### 3.5 *Arh3*-KO and -HT mice showed increased *in vivo* myocardial ischemia- reperfusion injury, which was significantly reduced by rucaparib

To investigate the role of ARH3 in cardiac protection *in vivo*, we investigated the effect of ARH3 on *in vivo* myocardial injury using IR. Myocardial infarction *in vivo* replicated the IR injury seen with the Langendorff-perfused isolated heart model. The infarct size in the *Arh3*- KO (male, 32.3 ± 1.8%, *n* = 8; female, 29.8 ± 1.2%, *n* = 8) mice was significantly larger than that of the WT mice (male, 21.2 ± 0.8%, *n* = 13; female, 19.0 ± 1.3%, *n* = 7) after their hearts were subjected to 2 h of ischemia, followed by 24 h of reperfusion in both the male and female mice (*Figure 3E and F*). Next, we examined the effect of the FDA- approved PARP inhibitor, rucaparib, on *in vivo* myocardial injury. Mouse hearts were subjected to 90 min of ischemia and 24 h of reperfusion with or without DMSO or 10 mg/kg rucaparib. Drug or control injection time courses are shown in *Figure 3G*. The infarct size of the *Arh3*- KO hearts (26.8 ± 1.53%, *n* = 6) and the *Arh3*-HT hearts (29.3 ± 1.72%, *n* = 5) were significantly larger than those of WT hearts (12.3 ±1.54%, *n* = 5) after IR injury (*Figure 3H and I*). Furthermore, 10 mg/kg rucaparib significantly decreased the infarct size of *Arh3*-KO hearts (3.6± 0.49%, *n* = 3) compared with the non-treated *Arh3*-KO mice (26.8 ± 1.53%, *n* = 6) during IR injury (*Figure 3H and J*). These results suggest that *Arh3*-KO and -HT mice experienced enhanced IR-induced myocardial injury. In addition, the FDA-approved PARP inhibitor rucaparib reduced the infarct size of mouse hearts with *Arh3* deficiency *in vivo* IR resulting from injury.

### 3.6 *Arh3*-KO and -HT C2C12 myoblasts and myotubes exhibited increased H2O2-induced cell death, which was rescued by transfection with the *Arh3* gene

We showed that *Arh3*-KO and -HT mice displayed reduced cardiac function and enhanced sensitivity to myocardial IR injury. To study the cellular consequences of *ARH3*-homozygous and -heterozygous mutations, we used mouse skeletal muscle cells (C2C12 myoblasts and myotubes) of each genotype. *Arh3*-WT, - HT, and -KO C2C12 cells were generated using CRISPR/Cas9 and ARH3-overexpressing C2C12 cell lines were generated by transfection with ARH3 expression vector (WT + ARH3 and KO + ARH3) to investigate the effect of ARH3 protein in cells. Endogenous ARH3 protein was identified in WT C2C12 myoblasts/myotubes and WT C2C12 myoblasts transformed with empty vector (EV) using Western blot analysis (*Figure 4A and B*). ARH3 protein was detectable in ARH3-overexpressing C2C12 myoblasts (WT + ARH3 and KO + ARH3) (*Figure 4B*). In contrast, ARH3 protein was expressed at lower levels relative to WT cells in HT C2C12 myoblasts/myotubes, but ARH3 was not expressed in KO cell lines (*Figure 4A and B*). Cytotoxicity assay following hydrogen peroxide (H2O2) exposure (0.1-1 mM) for 3 h of C2C12 myoblasts showed that viability of KO and HT myoblasts was significantly less than that of WT myoblasts (*Figure 4C*, upper left). Gabor Oláh *et al*.^22^ observed that when myoblasts were induced to differentiate to myotubes, PARP expression and PAR level were reduced. Concurrently, cells became more resistant to H2O2-induced oxidative stress.^22^ We therefore examined myotubes by cytotoxicity assay using high concentrations of H2O2 (0.5-10 mM) to examine difference in viability among genotypes.

**Figure 4.**
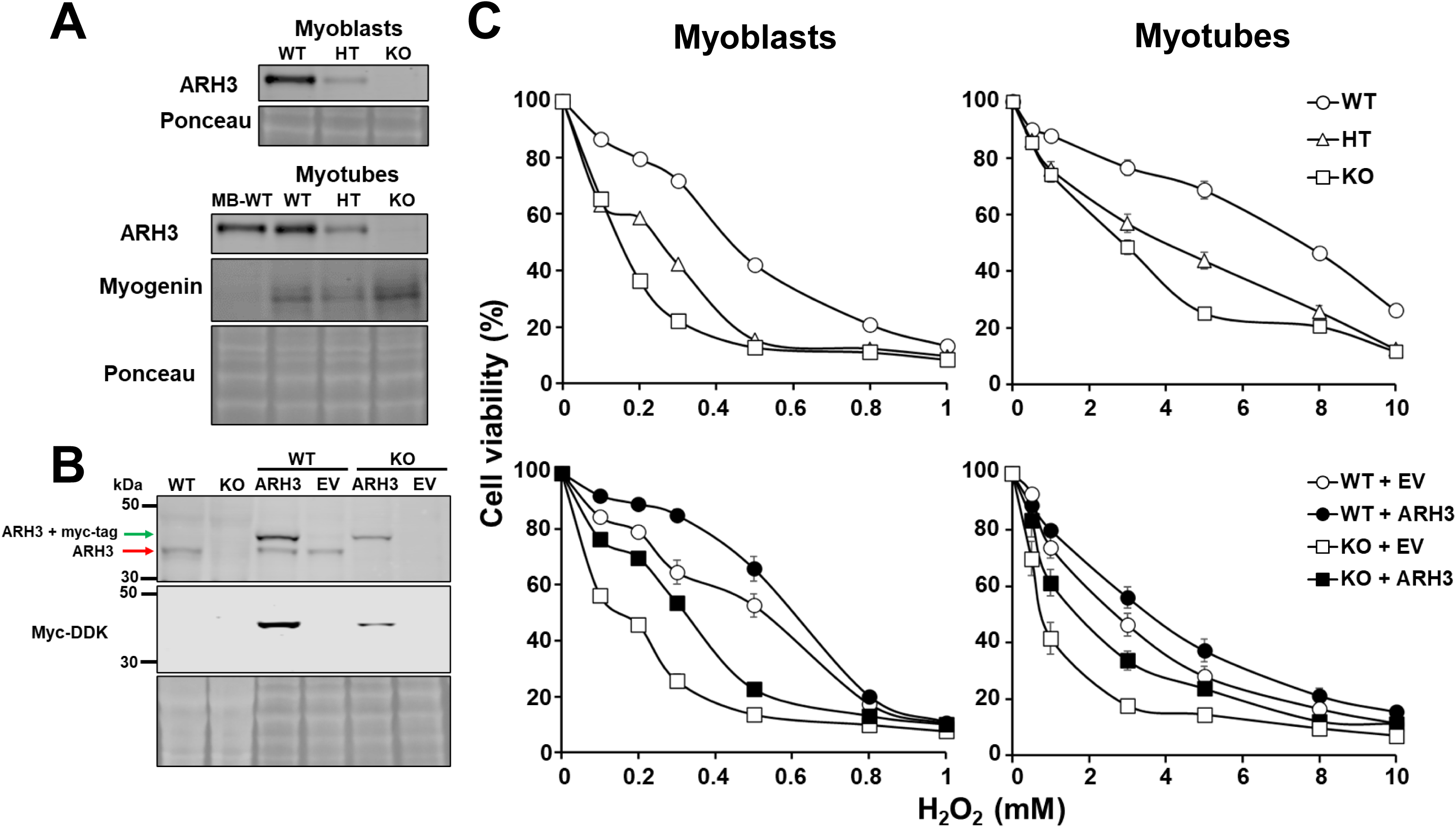
*Arh3*-KO and -HT C2C12 myoblasts and myotubes displayed enhanced H2O2-induced cell death, which was rescued by transfection with the *Arh3* gene. (*A*) ARH3 protein levels in CRISPR/Cas9-induced ARH3 mutants, corresponding to the WT, HT, and KO in C2C12 myoblasts (MB) and myotubes. Cells were subjected to Western blotting by using anti-ARH3 antibody or anti-myogenin antibody as a differentiation marker. Ponceau S staining shows whole lysates as an internal control. (*B*) C2C12 myoblasts overexpressing ARH3. Cells were subjected to Western blotting using anti-ARH3 antibody or anti-myc-DDK-tag antibody. Ponceau S staining shows whole lysates as an internal control. Empty vector (EV) was used instead of ARH3 expression vector as a negative control. CRISPR/Cas9-mediated, *Arh3-*edited C2C12 myoblasts were used as WT or KO cells. (*C*) Cell viability of C2C12 myoblast, myotubes, and ARH3-overexpressing C2C12 myoblasts and myotubes following H2O2 exposure. Cells were exposed to H2O2 (3 h) at indicated concentrations (means ± SEM, *n* = 6). Nonlinear regression analysis, WT vs HT, WT vs KO, WT + EV vs KO + EV, or KO + EV vs KO + ARH3 C2C12 myoblasts with *p* < 0.0001; WT + EV vs WT + ARH3 C2C12 myoblasts with *p* = 0.0033; WT vs HT, WT vs KO, or WT + EV vs KO + EV C2C12 myotubes with *p* < 0.0001; KO + EV vs KO + ARH3 C2C12 myotubes with *p* = 0.0089.

Consistent with the results seen in myoblasts, viability of KO and HT myotubes following H2O2 exposure (0.5-10 mM) for 3 h was significantly less than that of WT myotubes (*Figure 4C*, upper right). Previously, we showed that *Arh3*-KO mouse embryonic fibroblasts (MEFs), but not *Arh3*-HT MEFs, exhibited decreased viability following H2O2 exposure, resulting in PAR-dependent cell death or parthanatos.^18^ To determine the effect of heterozygosity in the C2C12 study, we performed cytotoxicity assays with H2O2 using two different *Arh3*-HT MEFs as a control. Endogenous ARH3 protein was identified in WT MEFs (*Figure S4A*). As expected, ARH3 protein was expressed at lower levels relative to WT cells in two different HT MEFs (*Figure S4A*). Consistent with the observations following exposure of C2C12 myoblasts/myotubes to H2O2, following H2O2 exposure (0.1-1 mM) for 3 h the viability of KO and two different HT MEFs was significantly less than that of the WT MEFs (*Figure S4B*). We previously reported that the two *Arh3*-HT MEFs were not susceptible to H2O2 (0.1-0.6 mM) following 24-h exposure.^18^ Since a half-life of H2O2 in culture medium is less than 15 min at concentrations between 0.6 to 6 mM,^23^ *Arh3*-HT MEFs may have sufficient PAR-hydrolase activity during long exposure, leading to recovery from H2O2-induced cell death, and thus, showing no difference in cytotoxicity between WT and HT MEFs.^18^ In conclusion, loss of *Arh3* in heterozygous and homozygous cells resulted in enhanced susceptibility to H2O2 in different murine cell lines, C2C12 myoblasts/myotubes, and MEFs. In addition, in *Arh3*-KO C2C12 myoblasts/myotubes transfected with the *Arh3* gene, viability of KO + ARH3 cells was significantly greater than that of KO + EV cells, following H2O2 exposure in both myoblasts and myotubes (*Figure 4C*, lower left and right). This result is consistent with our previous findings in MEFs transfected with the *Arh3* gene.^18^ In addition, the viability of WT + ARH3 myoblasts was significantly increased when compared to the WT + EV myoblasts following H2O2 exposure, but not in myotubes (*Figure 4C*, lower left and right). Thus, ARH3 exerts protective effects against H2O2-induced cell death in C2C12 myoblasts/myotubes.

### 3.7 Rucaparib blocked PAR-dependent cell death caused by H2O2-induced oxidative stress in C2C12 myoblasts and myotubes

We showed that ARH3 has a protective effect against H2O2- induced oxidative stress response in C2C12 myoblasts/myotubes. ARH3 is responsible for degradation of PAR, which should attenuate the oxidative stress response. To determine whether H2O2-induced cell death pathway is PAR-dependent, we monitored PAR levels and cytotoxicity following H2O2 exposure in the presence or absence of PARP inhibitors in C2C12 myoblasts/myotubes of different *Arh3* genotypes. Indicated concentrations of rucaparib or olaparib given for 30 min before 0.5 mM H2O2 exposure for 3 h significantly inhibited death of C2C12 myoblasts in a concentration-dependent manner (rucaparib: WT EC50 = 1.92 nM, KO EC50 = 341.2 nM, *Figure 5A*; olaparib: WT EC50 = 1.90 nM, KO EC50 = 105.6 nM, *Figure S5A*). In addition, 10 μM rucaparib significantly inhibited H2O2-induced death of both myoblasts and myotubes, regardless of genotypes (*Figure 5B*). Furthermore, two different PARP inhibitors, e.g., PJ34 (10 μM), olaparib (10 μM), significantly inhibited H2O2-induced death of myoblasts, but inhibitors of apoptosis (zVAD-fmk, 20 μM), necrosis (IM54, 10 μM), and necroptosis (necrostatin-1, 10 μM) did not have an effect (*Figure S5B*).

**Figure 5.**
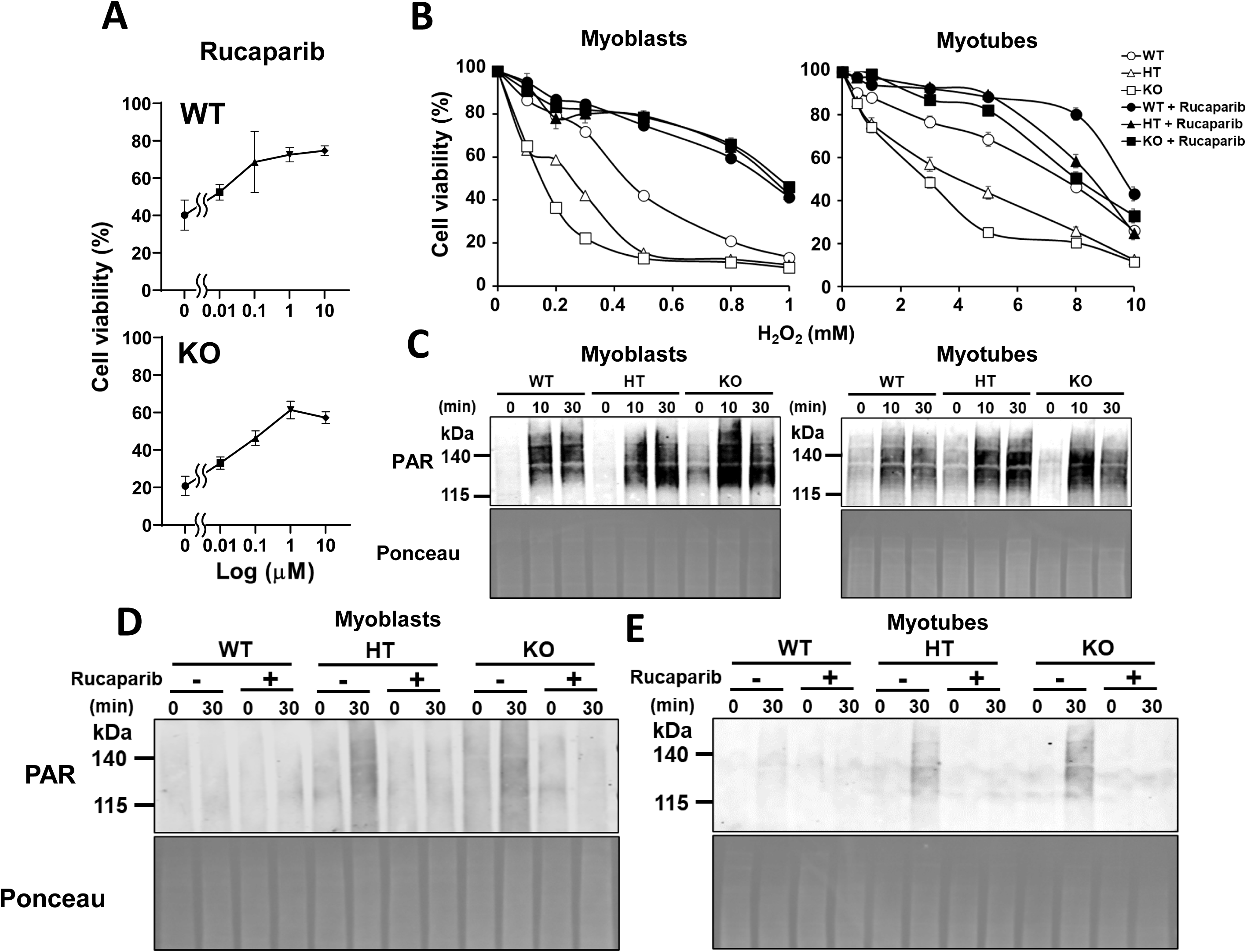
PARP inhibitors prevented PAR accumulation that was involved in H2O2-induced cell death pathway in C2C12 myoblasts and myotubes. (*A*) Concentration-response curve of rucaparib following H2O2 exposure. *Arh3*-WT or -KO C2C12 myoblasts were exposed to the indicated concentrations of rucaparib for 30 minutes before 0.5 mM H2O2 exposure for 3 h. Cell viability was calculated by comparison to the control group (100%; cell viability without H2O2). (WT: EC50 = 1.92 nM, KO: EC50 = 341.2 nM, means ± SEM, *n* = 3). (*B*) Cytotoxicity assay following H2O2 exposure with or without rucaparib (10 μM) between genotypes in myoblasts (Left) and myotubes (Right) (means ± SEM, *n* = 4). Nonlinear regression analysis, WT vs WT + rucaparib, HT vs HT + rucaparib, or KO vs KO + rucaparib C2C12 myoblasts/myotubes with *p* < 0.0001. (*C*) PAR levels and the amount of ARH3 protein in C2C12 myoblasts (Left) and myotubes (Left) between genotypes following 300 μM H2O2 exposure. Cells were subjected to Western blotting using anti- PAR (10H) antibody. Ponceau S staining shows whole lysates as an internal control. (*D*) Representative pictures of PAR levels between genotypes in myoblasts (Left) and myotubes (Right) with or without 10 μM rucaparib. Cells were subjected to Western blotting with anti- poly(ADP-ribose) (10H) antibody. Ponceau S staining shows whole lysates as an internal control.

In support of the effects on C2C12 myoblasts and myotubes with PARP inhibitors, 10 μM rucaparib or 10 μM olaparib significantly inhibited H2O2-induced death of MEFs (*Figure S4C and D*). Consistent with the results of cytotoxicity assays, the PAR levels in *Arh3*-KO and -HT C2C12 myoblasts and myotubes were higher than in the WT cells following 0.3 mM H2O2 exposure (*Figure 5C*). In addition, rucaparib reduced PAR levels in both myoblasts and myotubes (*Figure 5D and E*). Thus, PARP activation following H2O2 exposure induced PAR accumulation, resulting in PAR-dependent cell death, which is blocked by PARP inhibitors in myoblasts and myotubes.

## 4. Discussion

Based on our study, we show that (1) ARH3 is involved in cardiac function as well as protection under oxidative stress by controlling PAR levels of *Arh3* mutant mice (*Figure 1-3 and Figure S2 and 3*), (2) a PARP inhibitor, rucaparib, improves murine cardiac dysfunction (*Figure 2-3 and Figure S3*), (3) ARH3 confers cellular protection under oxidative stress by controlling PAR levels of C2C12 myoblasts and myotubes (*Figure 4, 5 and Figure S5*), (4) PARP inhibitors, e.g. rucaparib, olaparib, PJ34, prevented PAR-dependent cell death under oxidative stress in myoblasts/myotubes seen with *Arh3* mutations (*Figure 5 and Figure S5*).

Cardiac phenotypes, cardiac hypertrophy and a significant increase in heart weight/body weight ratio, were observed in *Arh3*-KO mice compared to WT mice (*Figure 1B and C*). Pathological cardiac hypertrophy has been characterized by cardiac fibrosis, increased cardiomyocyte size, and cardiac dysfunction.^24^ Although we found no evidence of cardiac fibrosis and cardiomyocyte disarray in *Arh3*-KO mouse hearts in the histological sections by Masson’s trichrome staining (data not shown), *Arh3*-KO mice showed significant increase in heart weight/body weight ratio, lower cardiac function by MRI, under dobutamine infusion by echocardiography, and by a Langendorff-perfused isolated heart model (*Figure 1D-H, 3A, and Figure S2*), suggesting that cardiac hypertrophy observed in *Arh3*-KO mice affected function.

Of interest, *Arh3*-HT mice showed results similar to those seen in *Arh3*- KO mice (*Figure 1H, 2B* and *Figure S2*). Our recent report showed that *Arh3*-KO mice displayed increased size of IR infarcts after brain IR injury, enhanced cytotoxicity following H2O2 oxidative stress, but WT and HT mice and cells (patient fibroblasts and MEFs), respectively, did not.^17^ In earlier reports, all patients had to have a bi-allelic *ARH3* homozygous mutation. Recently, *Lu et al.*^16^ identified a patient with a *ARH3*- heterozygous mutation with a neurodegenerative phenotype, suggesting that *ARH3* heterozygosity caused neurological symptoms. Similarly, our findings support the conclusion that *Arh3* heterozygosity in mice results in cardiac dysfunction (*Figure 1H, 2B, 3H,* and *I*).

In the treatment assay, the PARP inhibitor rucaparib improved cardiac contractility during a dobutamine stress test in all genotypes (*Figure 2*). In addition, rucaparib reduced the size of myocardial IR infarcts in *Arh3*- KO mice (*Figure 3H, J*). In parallel research, we observed that *Arh3*-KO and -HT C2C12 myoblasts and myotubes showed enhanced susceptibility to H2O2, leading to PARP activation, and PAR-dependent cell death, which was reduced by PARP inhibitors (*Figure 5* and *Figure S5*). Thus, PARP inhibitors might control cardiac abnormalities in patients with *ARH3* homozygous as well as heterozygous mutations.

Prior studies of patients with *ARH3* deficiency attributed their cardiac abnormalities to autonomic nervous system dysfunction. However, our findings on IR injury using the *ex vivo* Langendorff-perfused isolated mouse heart model showed that *Arh3*-KO mouse hearts displayed (1) lower cardiac function as assessed by the heart function recovery rate (post-ischemic rate pressure products), (2) enhanced size of the IR infarcts, and (3) increased IR-induced PAR levels compared to the WT mouse hearts (*Figure 3A-D*). Thus, ARH3 plays a direct role in cardiac function and protection by controlling PAR metabolism under IR injury. Consistent with the result seen *ex vivo* IR injury, *in vivo* IR injury enhanced size of the infarction in *Arh3*-KO and -HT mice (*Figure 3E, F, H,* and *I*), suggesting that ARH3 plays a critical role in cardiac protection under oxidative stress *in vivo*.

Our findings using rucaparib and olaparib, clinical PARP inhibitors, support their beneficial effect against oxidative stress in *in vitro* and *in vivo* studies. PARP inhibitors, which inhibit the catalytic activity of PARP and trap PARP on damaged DNA sites, are effective anti-cancer drugs.^22, 25–27^ PARP inhibitors—rucaparib, olaparib, niraparib, and talazoparib—have been approved for the treatment of ovarian, breast and pancreatic cancer with mutation in the homologous recombination repair genes BRCA-1/2, resulting in synthetic lethality.^22, 25, 26^ For treatment of other diseases candidates, classical PARP inhibitors may reduce cardiac dysfunction by improving cardiac contractility and reducing the size of cardiac infarction by myocardial IR injury in experimental animal models.^3, 19–21^ Although PARP inhibitors reduced cardiac dysfunction, PARP inhibitors are not well studied in the heart. Rucaparib and olaparib exhibit high-binding affinity for PARP-1, which is responsible for approximately 90% of PAR formation (IC50 rucaparib: 0.8-3.2 nM, IC50 olaparib: 1-19 nM; values were obtained from ChEMBL database).^28^ Our findings suggest that those PARP inhibitors show pharmacological benefit in treating cardiac dysfunction (*Figure 2 and 3J*).

### 4.1 Limitations

We found that *Arh3* contributes to cardiac function and protection directly in the murine heart. However, since the *Arh3* mutant mice showed neurological dysfunction assessed by motor activities (e.g., open-field, rotarod tests, isometric torque tests) (*Figure S1*), we need further investigation whether *Arh3* mutations might also affect the autonomic nervous system, thereby inducing cardiac dysfunction. The sympathetic autonomic nervous system appears to play an important role in the brain-heart interactions after stroke.^29–31^ In a prior report, brain IR injury reduced cardiac contractility, enhanced myocardial vulnerability to IR injury caused by altered sympathetic hyperactivity, increased nitro-oxidative signaling, and decreased short-term cardiac expression of JAK2, STAT3, and phosphorylated STAT3 in protein.^30, 31^ To determine if autonomic function is associated with the development of cardiac dysfunctions in *Arh3*-KO mice, evaluation of cardiac function by echocardiography and the levels of sympathetic stimulations (e.g., catecholamine, cortisol) in mouse heart tissue after brain IR injury *in vivo* and *ex vivo* are further needed.

## Supporting information

MRI video of ARH3 mice

## 5. Summary

In conclusion, we have presented the importance of the action of ARH3 in myocardium through mouse and cellular models with an FDA-approved PARP inhibitor. Our findings show that ARH3 controls PAR levels in myocardium. Our in-depth understanding of the role of ARH3 can provide therapeutic information in treating cardiac abnormalities in patients who are *ARH3* knockouts on show heterozygousity. PARP inhibitors might offer beneficial effects in treating the cardiac dysfunction in patients with *ARH3* deficiency.

## Supplemental material

Supplementary material is available at *Cardiovascular Research* online.

## Authors’ contributions

S.Y. and J.M. designed experiments. J.K., H.I.E., and X.B. helped to design experiments. J.K. and J.M. prepared animal study protocols. S.Y., J.K., H.I.E., D.S., and M.P. performed behavioral experiments. D.S, and A.N. performed echocardiography experiments. M.L. performed MRI experiments. X.B. performed *ex vivo* IR injury and Western blotting of the heart. H.S., R.C., and K.K performed *in vivo* IR injury. C.L. and F.Z. designed and constructed *Arh3*-KO and -HT C2C12 cells. Z.Y. prepared histological sections. S.Y. prepared samples and performed cytotoxicity assay and Western blotting experiments. S.Y. and J.M. wrote the manuscript. All authors reviewed the manuscript.

## Funding

This work was supported by the Intramural Research Program, National Institutes of Health, National Heart, Lung, and Blood Institute [grant number: ZIA-HL-000659].

## Acknowledgements

We are thankful to Dr. Rodney L. Levine (Senior Investigator, NHLBI) for helpful advice and discussions.

## Conflict of interest

None declared.

## Data availability

The data relating to this article are available in the article itself or in its online Supplementary material.

## Supplementary Material

### Supplementary Methods

#### Generation of *Arh3*-KO and -HT mice

Generation of *Arh3-*KO and -HT mice was described by Mashimo et al.^1^ *Arh3-*KO and - HT mice were backcrossed 9 times with C57BL6/L mice.^1^ Genotypes of these mice were confirmed by PCR.^1^

#### Open-field test

General locomotor activity was examined in 3-4- and 7-8-month-old mice, using the open-field test. Mice were individually placed in a 15” X 15” X 15” Perspex arena viewing chamber and allowed to freely explore for 30 min. Recordings of the test were analyzed using video behavioral tracking software (ANY-maze, Stoelting Co.) and edge time, distance traveled, time immobile and time mobile, and velocity of movement was measured in 5-min to 30-min time bins.

#### Rotarod test

Three-four- or 7-8-month-old mice were subjected to rotarod test. This test was used to assess motor coordination, balance, and equilibrium using a Rotamex 5 rotarod apparatus (Columbus Instruments, Columbus, OH). The day before the test, mice received two acclimation sessions to the rotarod at low constant speeds (static, 4rpm) with 3 one-min trials for the static rod and 3 three-minute trials/training for the 4rpm, with 10-min rest intervals between trials and 2 h between sessions. On test day, the rotarod was set at an accelerating mode of 4-40 rpm for 5 min and the latencies for the mice to fall off of the rotating rod were recorded. The test was repeated three times with 1-h rest between each trial.

#### Isometric torque test

Ten-month old mice were anesthetized with 2-3% isoflurane and maintained at 37°C. The hindlimb skin was shaved, disinfected, and then stabilized by a U-shaped holder to keep the leg in place during the test. Transcutaneous electrodes were inserted to stimulate hindlimb plantar flexor muscle contraction and the maximal isometric torque was assessed by an Aurora Scientific Inc model 1300A. A forced frequency protocol was performed first by administering an increasing progression of stimulation frequencies (25- 250 Hz in 25 Hz increments delivered once every minute) and the maximal isometric torque at peak tetanic contraction was recorded for each stimulation frequency. A fatigue protocol was then applied in which a series of 150 Hz stimulations were given every five seconds to assess fatigue response after 100 repeated contractions. Isometric torque was normalized to muscle mass and reported as specific torque.

#### Dobutamine stress with echocardiography

Dobutamine stress testing was performed using 10-month-old mice. Mice were placed on a heated imaging platform with electrocardiography leads and lightly anesthetized with 2% isoflurane via a nose cone during echocardiography and dobutamine infusions. Dobutamine was administered by intravenous infusion via a tail-vein catheter. After the baseline-scan, the mice received constant rate infusions of dobutamine (0.625 mg/mL in normal saline containing 5% dextrose). The first administration, a low dose, was 10 mg/kg/min. After the heart rate reached a steady state approximately 500 beats/min, a higher dose (40 mg/kg/min) was given. The length of time of the infusions was between approximately 20 and 30 min. Cardiac images were obtained using the Vevo2100 ultrasound system (FUJIFILM VisualSonics Inc., Toronto, Canada) with a 30-MHz ultrasound probe at baseline and during the low and high dose dobutamine infusions. For the treatment assay, PARP inhibitor rucaparib (MedChemExpress) at 10 mg/kg or vehicle buffer (1.25% Dimethyl sulfoxide (DMSO), 10% 2-hydroxypropyl-b-cyclodextrin (Sigma-Aldrich), 0.4% sodium chloride) was administered by intraperitoneal injection two times, 24 h and 1 h before dobutamine stress.

#### Reagents and antibodies

Rucaparib and olaparib were purchased from MedChemExpress; PJ34, pan-caspase inhibitor (zVAD-fmk), and mouse monoclonal anti-poly(ADP-ribose) (PAR) (10H) antibodies from Enzo Life Sciences; necrosis inhibitor (IM-54) from Santa-Cruz; necroptosis inhibitor (Necrostatin-1) from Millipore Sigma; mouse monoclonal anti-DDK antibodies from Origene; mouse monoclonal anti-myogenin antibodies from BD Biosciences; rabbit polyclonal anti-mouse ADP-ribosyl-acceptor hydrolase (ARH3) antibodies were produced in rabbit by injecting ARH3 peptide (YenZym antibodies LLC).

#### Cells

C2C12 cells and MEFs were cultured in Dulbecco’s Modified Eagle Medium (DMEM) supplemented with 10% heat-inactivated fetal bovine serum (FBS), penicillin (100 U/ml) and streptomycin (100 μg/ml) in a humidified incubator with 5% CO_2_ at 37.0°C. The *Arh3*- KO C2C12 cells were generated by CRISPR/Cas9 KO plasmids consisting of *Arh3*- specific 20 nt guide RNA sequences derived from the GeCKO (v2) library (Santa Cruz, sc-430308). The *Arh3*-KO and -HT MEFs were generated from WT and *Arh3*-KO littermates.^2^

#### Using CRISPR/Cas9 to generate C2C12-*Arh3* null and haploinsufficient mutants

The *Arh3* KO and HT mutant C2C12 cell lines were generated using CRISPR/Cas9. Briefly, two guide RNAs were designed using CHOPCHOP (https://chopchop.cbu.uib.no), one (*Arh3*-up: GCAACCTCGGAAGCGCGAGA) cutting shortly after the translation initiation codon (ATG) in Exon 1 and the other one (*Arh3*-down: GTAAGGGCCGCAGCTTCGGT) cutting near the stop codon in Exon 6 of the mouse *Arh3* gene. These crRNAs were purchased from Integrated DNA Technologies (IDT) and annealed with ATTO-550-labeled tracrRNA (IDT #1075928) to form complete guide RNAs, which were then co-incubated with Cas9 nuclease (IDT #1081059) at an equimolar ratio at room temp for 20 min to allow the formation of ribonucleotide-protein complexes (RNPs). The RNPs were electroporated into C2C12 cell using the Lonza 4D-Nucleofector X Unit. Briefly, ∼5 x 10^5^ cells were first mixed with the RNPs (4.7 µM final concentration) in nucleofection solution (Lonza Kit#VXC-2032) and then electroporated using Lonza’s pre-set program for C2C12 cells (CD-137). After the transfection, cells were immediately transferred into a well of a 24-well plate containing 0.5 ml prewarmed DMEM/10% FBS medium and incubated at 37°C, 5% CO_2_ for 24 hrs. Following recovery, the transfected cells were serial diluted and dispensed into 96-well plates for single-cell cloning. Those clones which grew were genotyped by PCR analyses of extracted genomic DNAs. Primer pair (Ex1-fw: TTTACAAATGAGAAGGTAGG; Ex1-rv: CTCTGAATGTGGTCATGGCTGG) were used for detecting small deletions around the Exon 1 cutting site, while primer pair (Ex6-fw: CCAGAGGACTCTCATCTACTCC; and Ex6-rv: CAGGCTGTCTCAAGATTCCAGC) were used for detecting small deletions near the Exon 6 cutting site. Primer pair Ex1-fw and Ex6-rv were used to identify clones with the entire coding region deleted. C2C12 clones with homozygous (-/-) and heterozygous Arh3 (+/-) deletions were grown and analyzed by Sanger sequencing for confirming desired deletions.

#### Cytotoxicity assays

C2C12 myoblasts and MEFs (1.0 × 10^4^ or 3 × 10^3^ cells) were incubated on 96-well plates for 1 day before exposure for 3 h to the indicated concentrations (0.1-1 mM) of H_2_O_2_. C2C12 myotubes (1.2 × 10^4^ cells) were incubated on 96-well plates for 1 day and then, media was changed to 2% horse serum and culture was continued for 5 days before exposure for 3 h to the indicated concentrations (0.1-1 mM) of H_2_O_2_. PARP inhibitors: rucaparib (0.1-10 μM), olaparib (0.1-10 μM), PJ34 (10 μM); Pan-caspase inhibitor: zVAD-fmk (20 μM); necrosis inhibitor: IM-54 (10 μM); necroptosis inhibitor: necstatin-1 (10 μM); were added for 30 min prior to H_2_O_2_ exposure. Cell viability was assessed using a Cell Counting Kit-8 (Dojindo) by measuring absorbance at 450 nm according to the manufacturer’s instructions. Absorbance was measured using SpectraMax M5 Microplate Reader (Molecular Devices).

#### SDS-PAGE and Western blotting

C2C12 myoblasts and MEFs (5.0 × 10^5^ cells) were incubated on 6-well plates for 1 day before exposure for 3 h to 0.3 mM of H_2_O_2_. Cells were lysed in 20 mM Tris-HCl (pH 7.4) with 2% SDS supplemented with a proteinase inhibitor (cOmplete^™^, EDTA-free Protease Inhibitor Cocktail, Roche). Protein concentration of lysates was measured using a bicinchoninic acid (BCA) assay kit (Thermo Scientific). Each sample was subjected to Bis-Tris SDS-PAGE (Invitrogen) and then transferred to nitrocellulose membranes (Invitrogen). Each membrane was blocked with Odyssey Blocking Buffer (LI-COR) or 2 % (w/v) Blocking-Grade Blocker (BIO-RAD) at room temp for 1 h and then incubated with primary antibody at 4°C overnight. Each membrane was washed with 1 x Tris-buffered saline containing 0.05% Tween 20 and then incubated with an appropriate secondary antibody (LI-COR) in blocking buffer for 1 h at room temperature. Each membrane was washed again with Tris-buffered saline and analyzed by an Odyssey Imaging Systems (LI-COR).

## Supplemental Figures and table

**Supplemental Figure 1.**
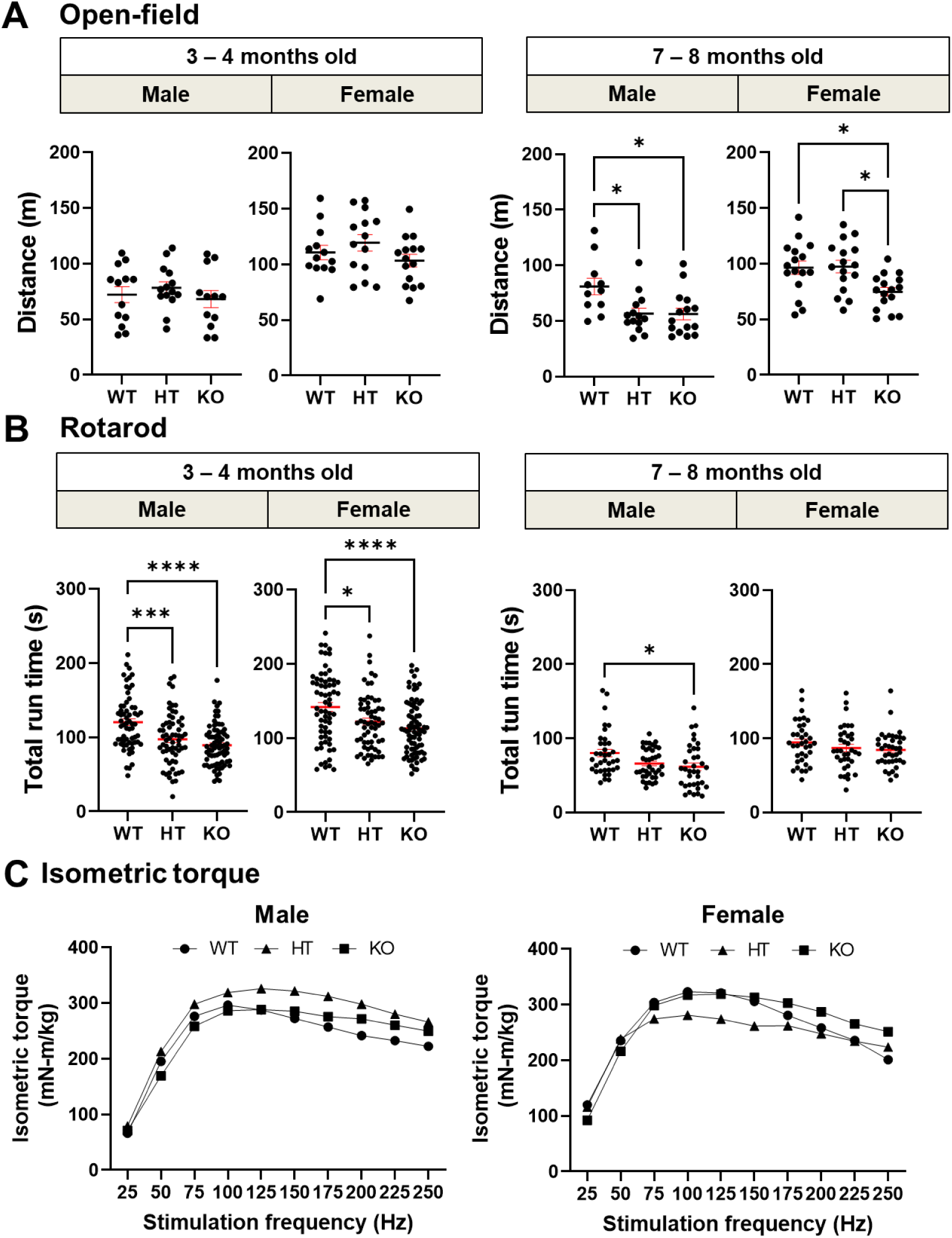
A*r*h3 mutant mice displayed impaired locomotor activity and motor coordination in an age- and gender-dependent manner assessed by open-field, rotarod, and isometric torque tests. (*A*) Locomotor activity assessed by open-field test in mice among *Arh3* genotypes. Two-way ANOVA Bonferroni’s multiple comparisons test, \**p* < 0.05, open-field test was performed three times in WT males, *n* = 13, 22.5, ± 2.5 weeks; HT males, *n* = 14, 22.3 ± 2.8 weeks; KO males, *n* = 12, 22.4 ± 2.5 weeks; WT females, *n* = 13, 26.2 ± 3.6 weeks; HT females, *n* = 14, 24.3 ± 0.7 weeks; KO females, *n* = 8, 25.3 ± 3.4 weeks (3-4 months); WT males, *n* = 11, 30.3, ± 1.7 weeks; HT males, *n* = 14, 29.8 ± 1.7 weeks; KO males, *n* = 15, 30.4 ± 2.0 weeks; WT female, *n* = 16, 29.9 ± 2.4 weeks; HT females, *n* = 16, 29.6 ± 2.9 weeks; KO females, *n* = 16, 29.4 ± 2.0 weeks (7-8 months). (*B*) Motor coordination assessed by rotarod test in mice among *Arh3* genotypes. Two-way ANOVA Bonferroni’s multiple comparisons test, \**p* < 0.05, \*\*\**p* < 0.001, \*\*\*\**p* < 0.0001, rotarod test was performed three times in WT males, *n* = 21, 25.7, ± 2.9 weeks; HT males, *n* = 21, 26.8 ± 3.6 weeks; KO males, *n* = 25, 26.8 ± 3.8 weeks; WT females, *n* = 22, 25.0 ± 2.0 weeks; HT females, *n* = 21, 26.5 ± 3.2 weeks; KO females, *n* = 26, 25.7 ± 3.5 weeks (3-4 months); WT males, *n* = 12, 34.0, ± 1.5 weeks; HT males, *n* = 13, 32.5 ± 1.3 weeks; KO males, *n* = 12, 32.7 ± 1.0 weeks; WT females, *n* = 12, 32.0 ± 1.4 weeks; HT females, *n* = 12, 32.0 ± 1.1 weeks; KO females, *n* = 13, 31.4 ± 1.3 weeks (7-8 months). (*C*) Muscle contractility assessed by isometric torque test in mice among *Arh3* genotypes. Isometric torque test was performed three times in WT males, *n* = 6, 38.6, ± 2.2 weeks; HT males, *n* = 6, 37.5 ± 2.2 weeks; KO males, *n* = 6, 37.0 ± 4.5 weeks; WT females, *n* = 6, 40.3 ± 3.0 weeks; HT females, *n* = 6, 39.1 ± 2.2 weeks; KO females, *n* = 6, 38.5 ± 3.0 weeks. Nonlinear regression analysis, there were no statistical differences among genotypes.

**Supplemental Table 1.** The performance of open-field and rotarod tests in mice among genotypes.

| Test | Gender | Mouse | Age (months) |  |
| --- | --- | --- | --- | --- |
|  |  |  | 3-4 | 7-8 |
| Open-field<br>(meter) | Male | WT | $72.1 \pm 1.99$ | $81.0 \pm 2.27$ |
| | | *Arh3*-HT | $78.2 \pm 1.42$ | $56.6 \pm 1.18^*$ |
| | | *Arh3*-KO | $68.1 \pm 2.21$ | $56.3 \pm 1.33^*$ |
| | Female | WT | $110.5 \pm 1.77$ | $96.5 \pm 1.49$ |
| | | *Arh3*-HT | $119.2 \pm 1.99$ | $97.4 \pm 1.41$ |
| | | *Arh3*-KO | $94.7 \pm 2.87$ | $74.8 \pm 0.99^*$ |
| Rotarod<br>(seconds) | Male | WT | $120.4 \pm 0.58$ | $80.0 \pm 0.85$ |
| | | *Arh3*-HT | $97.4 \pm 0.56^{***}$ | $66.0 \pm 0.52$ |
| | | *Arh3*-KO | $89.3 \pm 0.37^{****}$ | $61.7 \pm 0.84^*$ |
| | Female | WT | $141.6 \pm 0.72$ | $94.5 \pm 0.79$ |
| | | *Arh3*-HT | $122.1 \pm 0.60^*$ | $86.8 \pm 0.82$ |
| | | *Arh3*-KO | $113.0 \pm 0.46^{****}$ | $84.4 \pm 0.64$ |

Data represent mean ± SEM assessed by two-way ANOVA
Tukey's multiple comparison test. $*P < 0.05$ , $***P < 0.001$ , $****P < 0.0001$ .

**Supplemental Figure 2.**
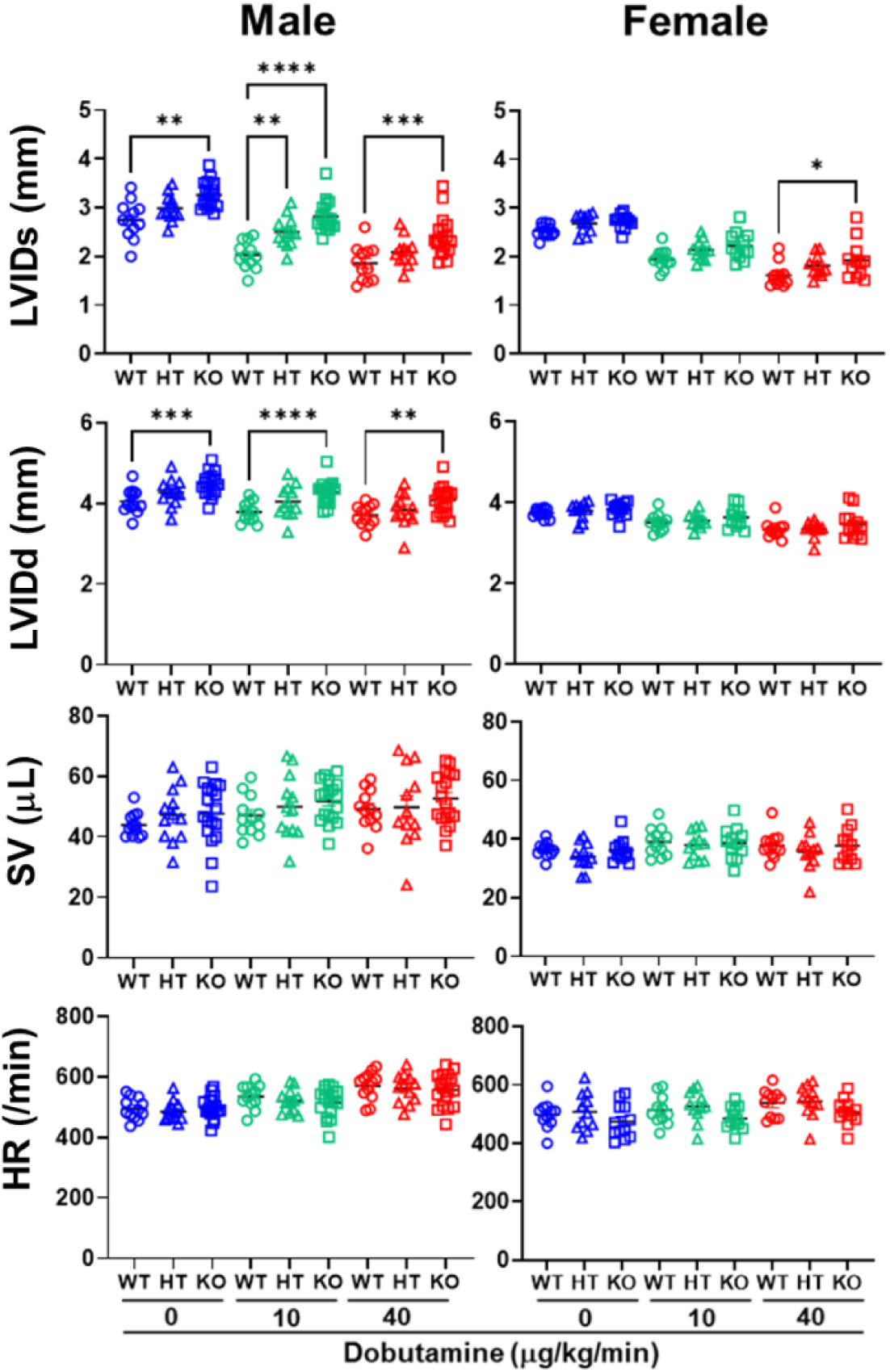
Function parameters among genotypes obtained by echocardiography. Assessment of left ventricular internal dimension (LVID) at systole (LVIDs), LVID at diastole (LVIDd), stroke volume (SV), and heart rate (HR) in male and female mice among genotypes subject to dobutamine stress (0, 10, and 40 mg/kg/min). Two-way ANOVA Tukey’s multiple comparisons test, \**p* < 0.05, \*\**p* < 0.01, \*\*\**p* < 0.001, \*\*\*\**p* < 0.0001, *n* = 8-12 (10 months).

**Supplemental Figure 3.**
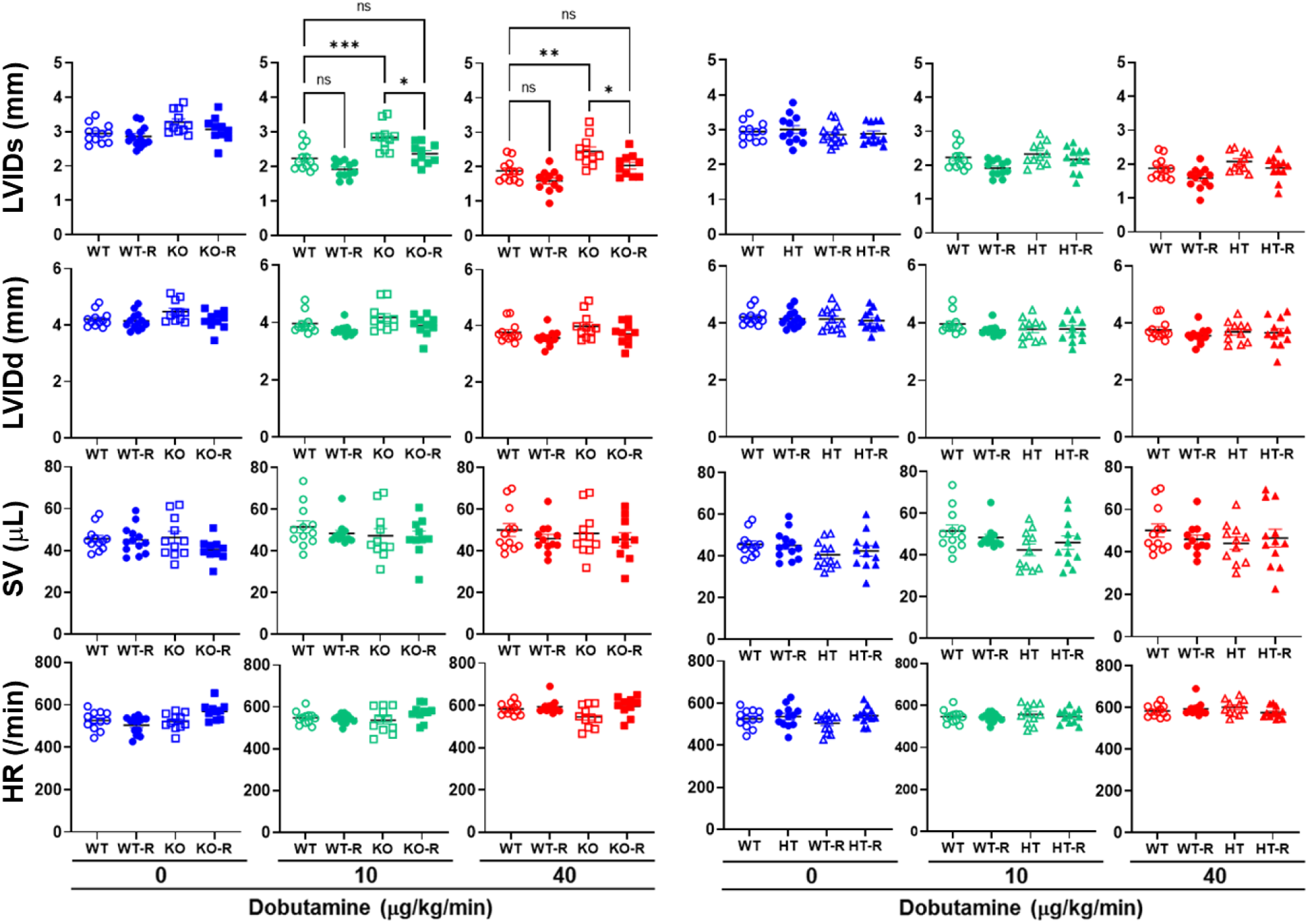
Cardiac function parameters among genotypes obtained by echocardiography with or without rucaparib. (*A*) Assessment of LVIDs, LVIDd, SV, and HR in male and female mice among genotypes subject to dobutamine stress (0, 10, and 40 mg/kg/min). Two-way ANOVA Tukey’s multiple comparisons test, ns: no significant difference, \**p* < 0.05, \*\**p* < 0.01, \*\*\**p* < 0.001, *n* = 8-12 (10 months).

**Supplemental Figure 4.**
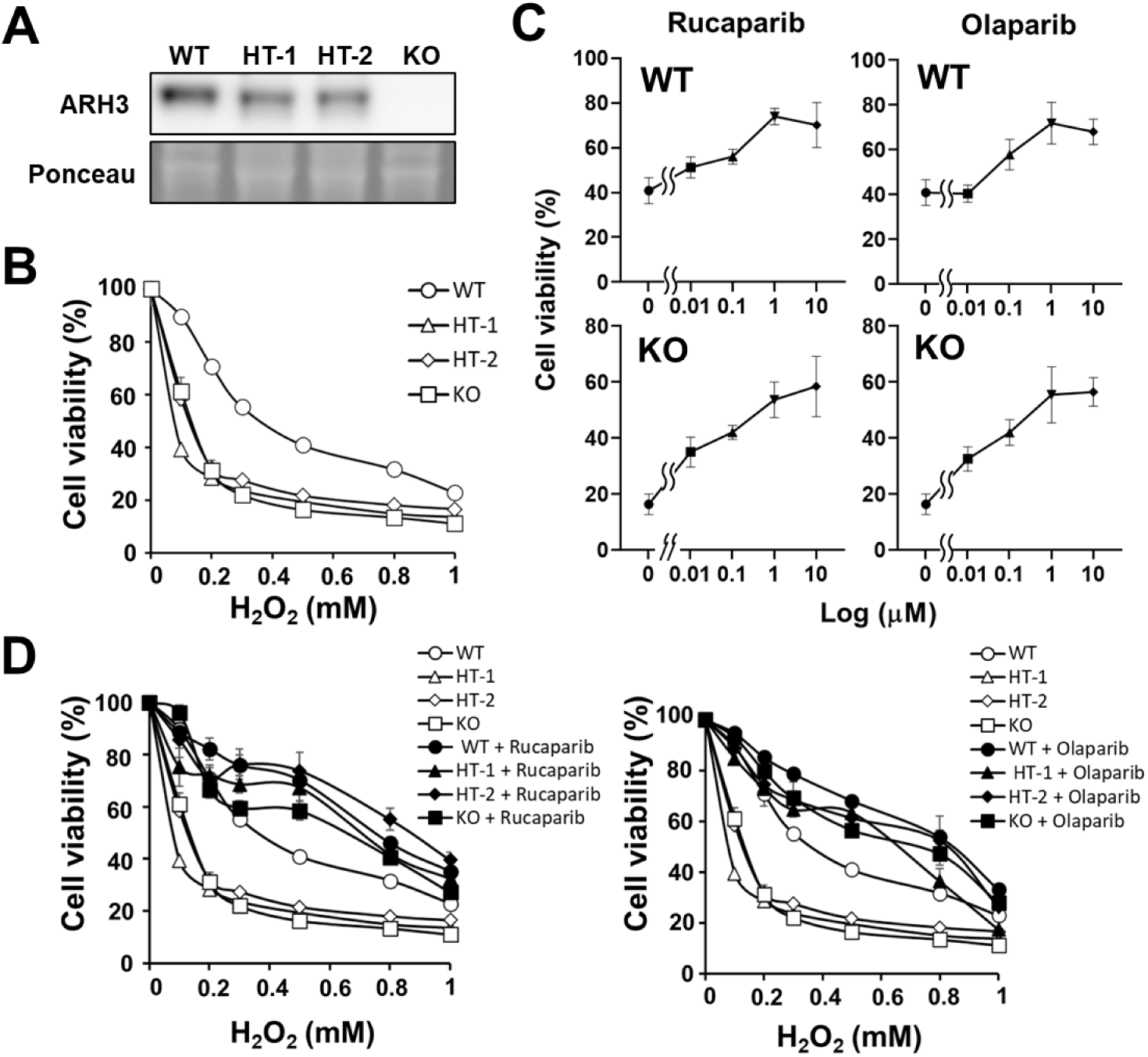
PARP inhibitors and ARH3 protected MEFs viability among genotypes following H_2_O_2_ exposure. (*A*) ARH3 expression of the *Arh3*-WT, HT, or KO MEFs. Cells were subjected to Western blotting with anti-ARH3 antibody. Ponceau S staining shows whole lysates as an internal control. (*B*) Cytotoxicity assay following H_2_O_2_ exposure with or without 10 μM rucaparib or 10 μM olaparib among MEFs genotypes. (means ± SEM, *n* = 4). Nonlinear regression analysis, WT vs HT-1, WT vs HT-2, WT vs KO MEFs with *p* < 0.0001. (*C*) Concentration-response of rucaparib or olaparib following H_2_O_2_ exposure. *Arh3*-WT or -KO MEFs were exposed to the indicated concentrations of rucaparib or olaparib for 30 min before 0.5 mM H_2_O_2_ exposure for 3 h. Cell viability was calculated by comparison to the control group (100%; cell viability without H_2_O_2_). (Rucaparib, WT: EC_50_ = 7.79 nM, KO: EC_50_ = 639.1 nM, means ± SEM, *n* = 3-16; Olaparib, WT: EC_50_ = 35.0 nM, KO: EC_50_ = 859.9 nM, means ± SEM, *n* = 4-16). (*D*) Cytotoxicity assay following H_2_O_2_ exposure with or without rucaparib (10 μM) (Left) or olaparib (10 μM) (Right) between genotypes in MEFs; (means ± SEM, *n* = 4). Nonlinear regression analysis, WT vs WT + rucaparib (or + olaparib), HT-1 vs HT-1 + rucaparib (or + olaparib), HT-2 vs HT-2 + rucaparib (or + olaparib), or KO vs KO + rucaparib (or + olaparib) MEFs with *p* < 0.0001.

**Supplemental Figure 5.**
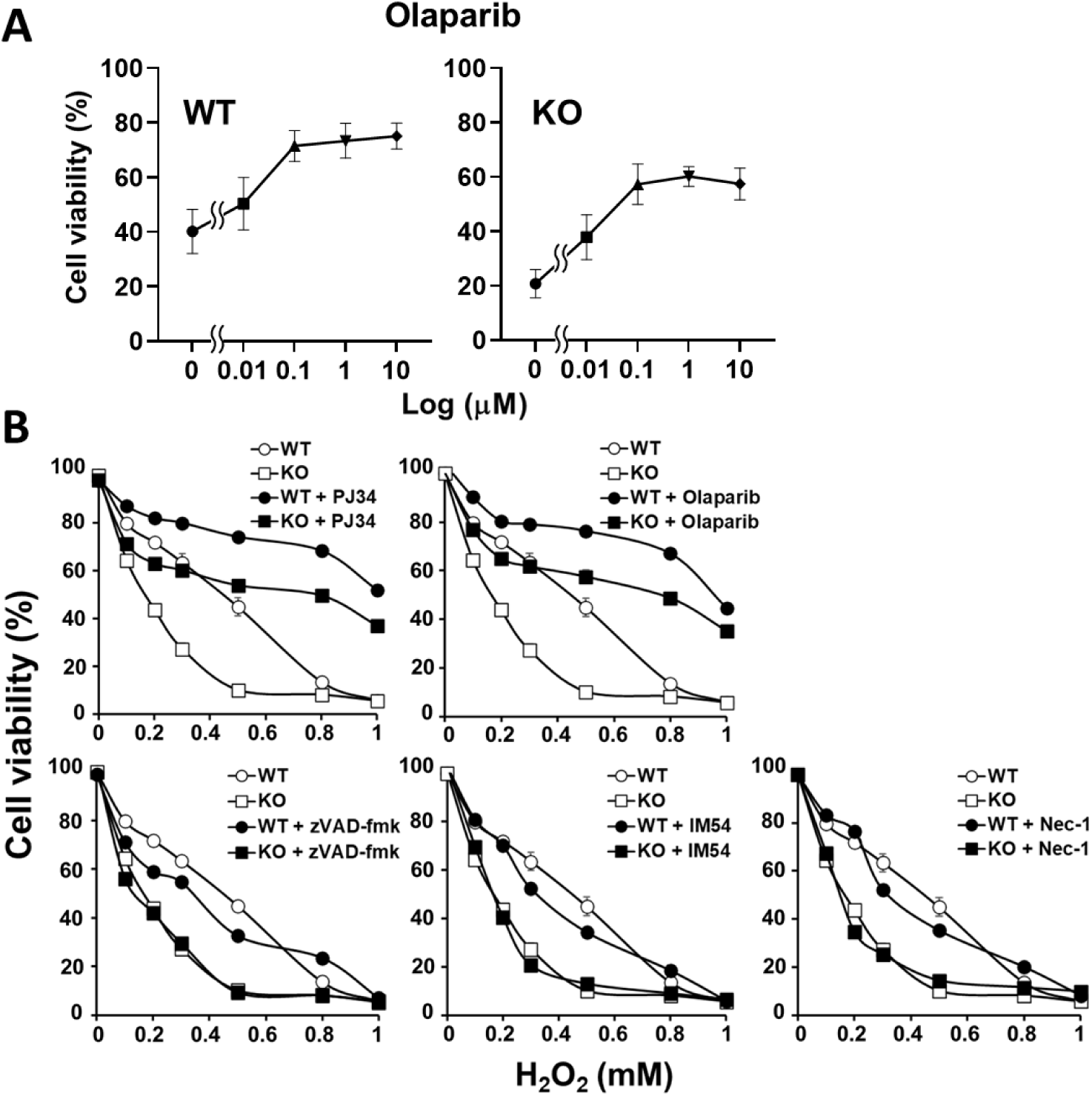
PARP inhibitors vs other inhibitors that participate in cell death pathways following H_2_O_2_ exposure of C2C12 myoblasts. (*A*) Concentration-response of rucaparib of olaparib following H_2_O_2_ exposure. *Arh3*-WT or -KO C2C12 myoblasts were exposed to the indicated concentrations of olaparib for 30 min before 0.5 mM H_2_O_2_ exposure for 3 h. Cell viability was calculated by comparison to the control group (100%; cell viability without H_2_O_2_). (WT: EC_50_ =1.9 nM, KO: EC_50_ = 105.6 nM, means ± SEM, *n* = 3). (*B*) Cytotoxicity assay following H_2_O_2_ exposure with or without PJ34 (10 μM), olaparib (10 μM), zVAD-fmk (20 μM), IM54 (10 μM), or necrostatin-1 (10 μM) between *Arh3*-WT or -KO C2C12 myoblasts. (means ± SEM, *n* = 4). Nonlinear regression analysis, WT vs KO, WT vs WT + olaparib (or + PJ34), or KO vs KO + olaparib (or + PJ34) C2C12 myoblasts with *p* < 0.0001.

## Full unedited gel images

Full unedited gel images for Figure 1A

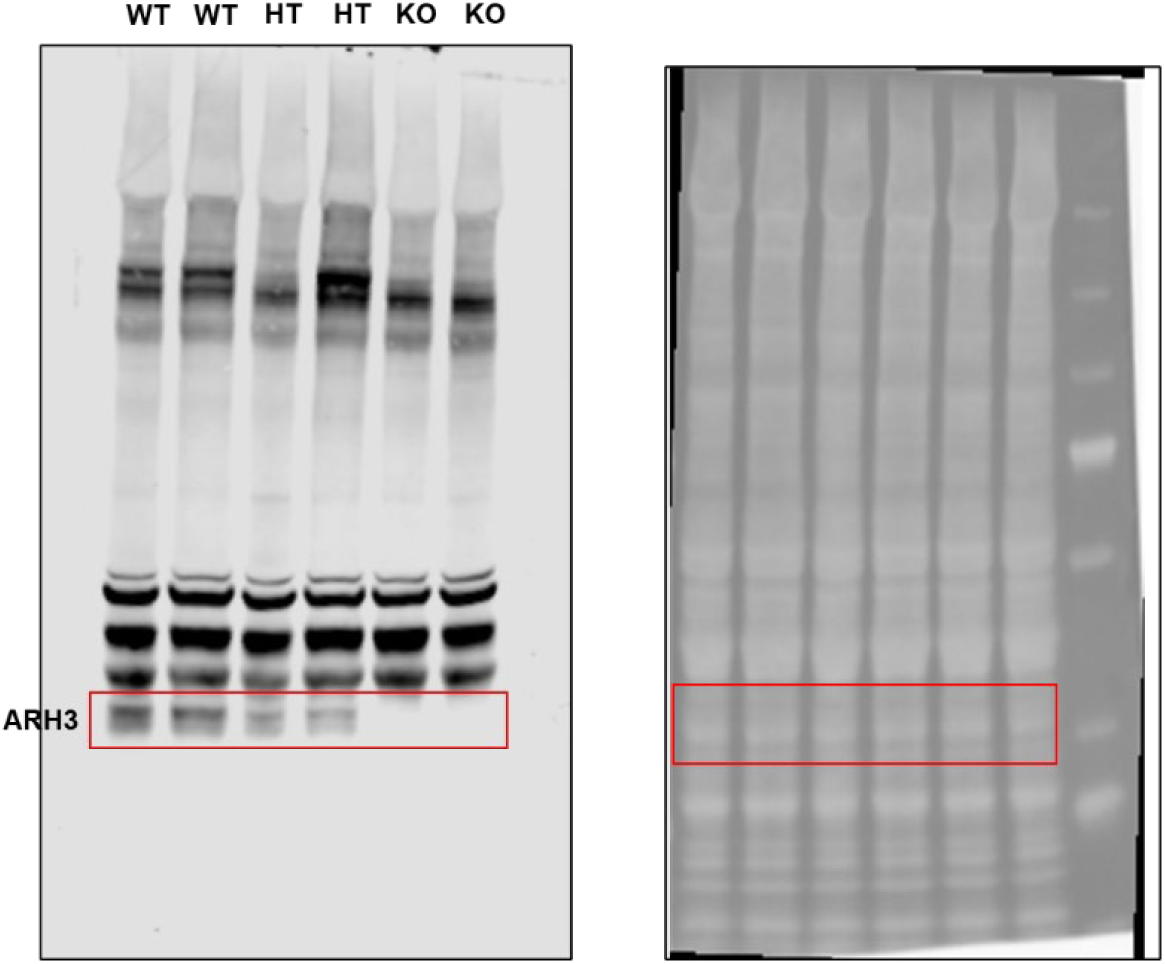

Full Unedited gel images for Figure 3D

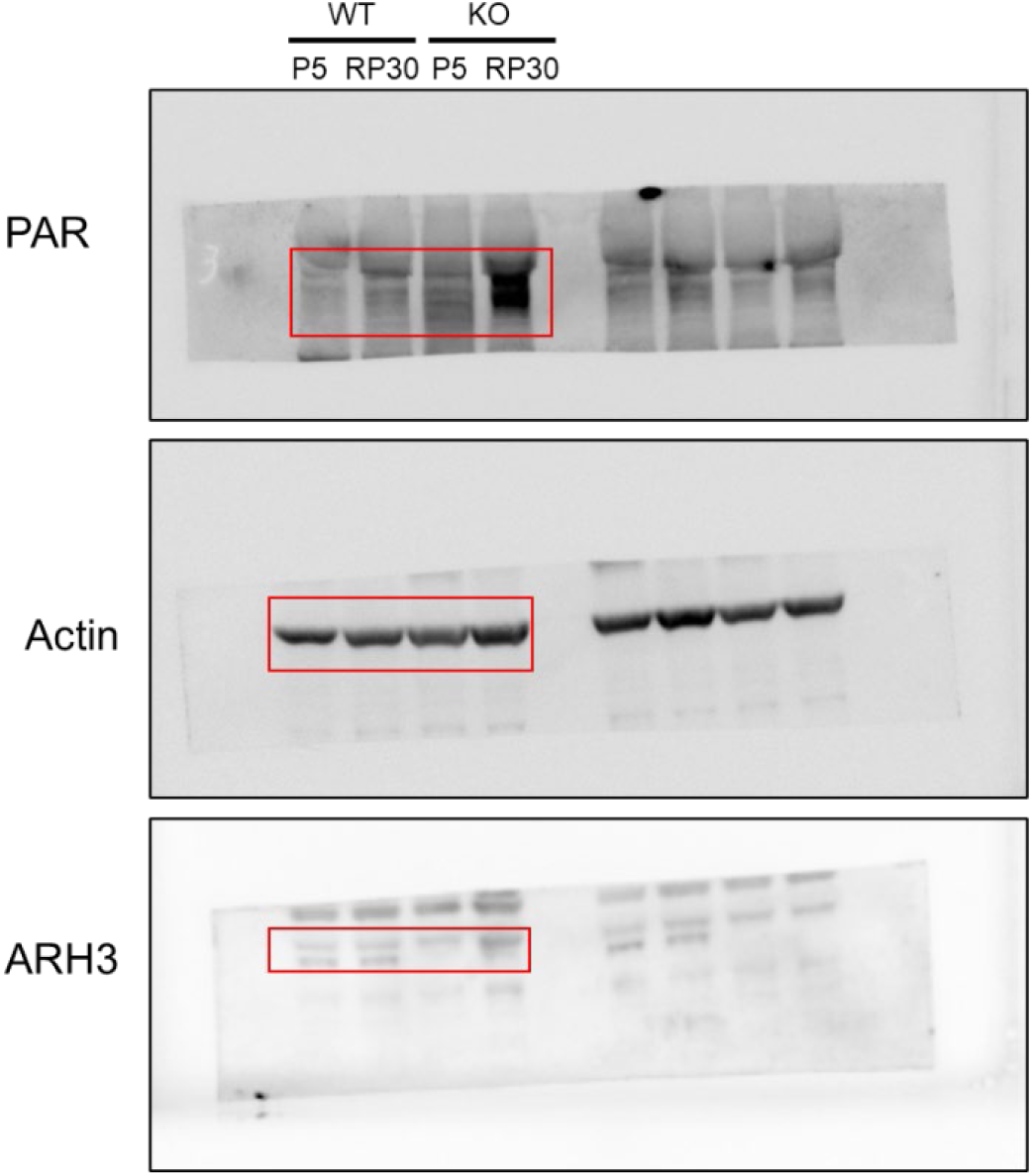

Full unedited gel images for Figure 4A

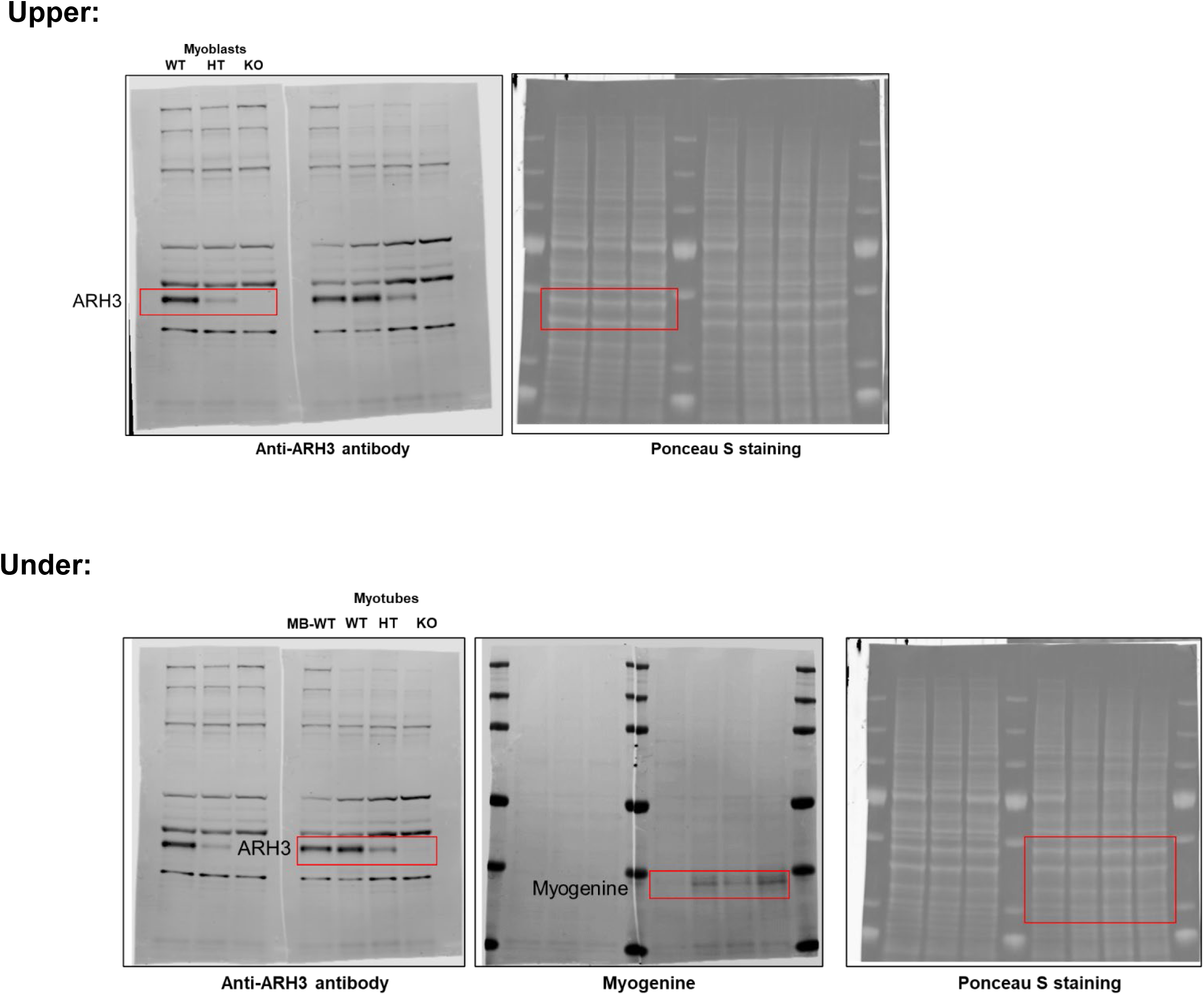

Full unedited gel images for Figure 4B

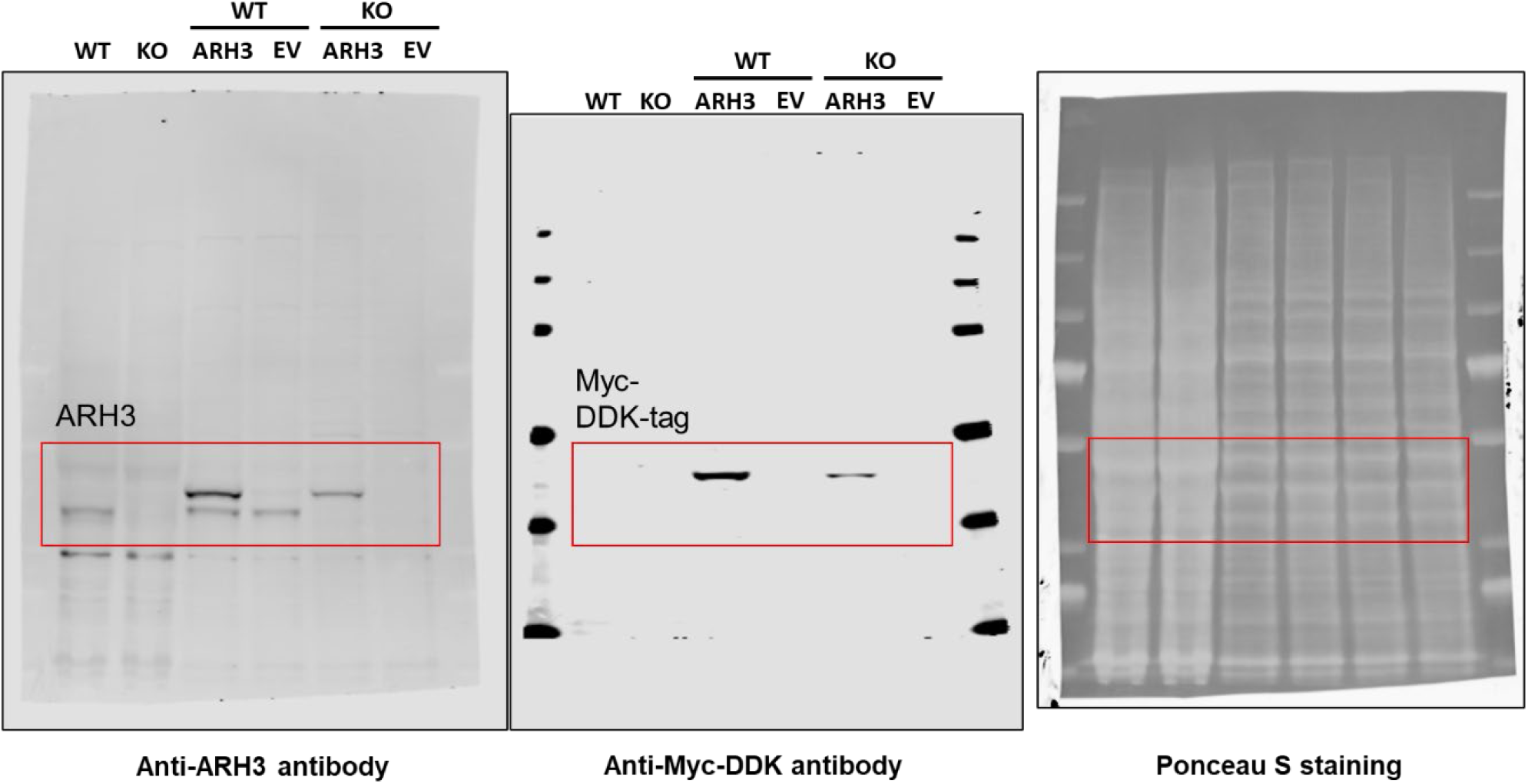

Full unedited gel images for Figure 5C

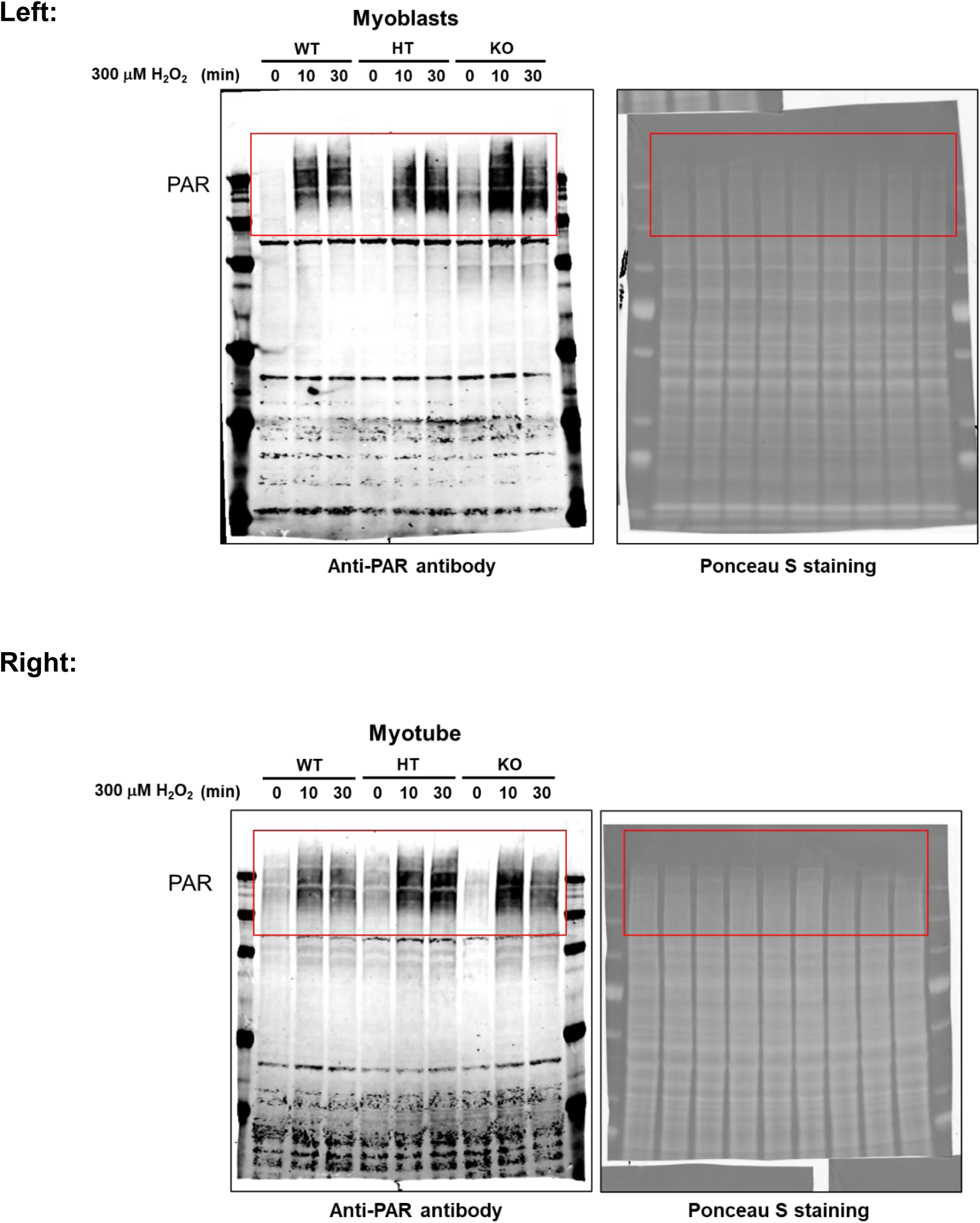

Full unedited gel images for Figure 5D

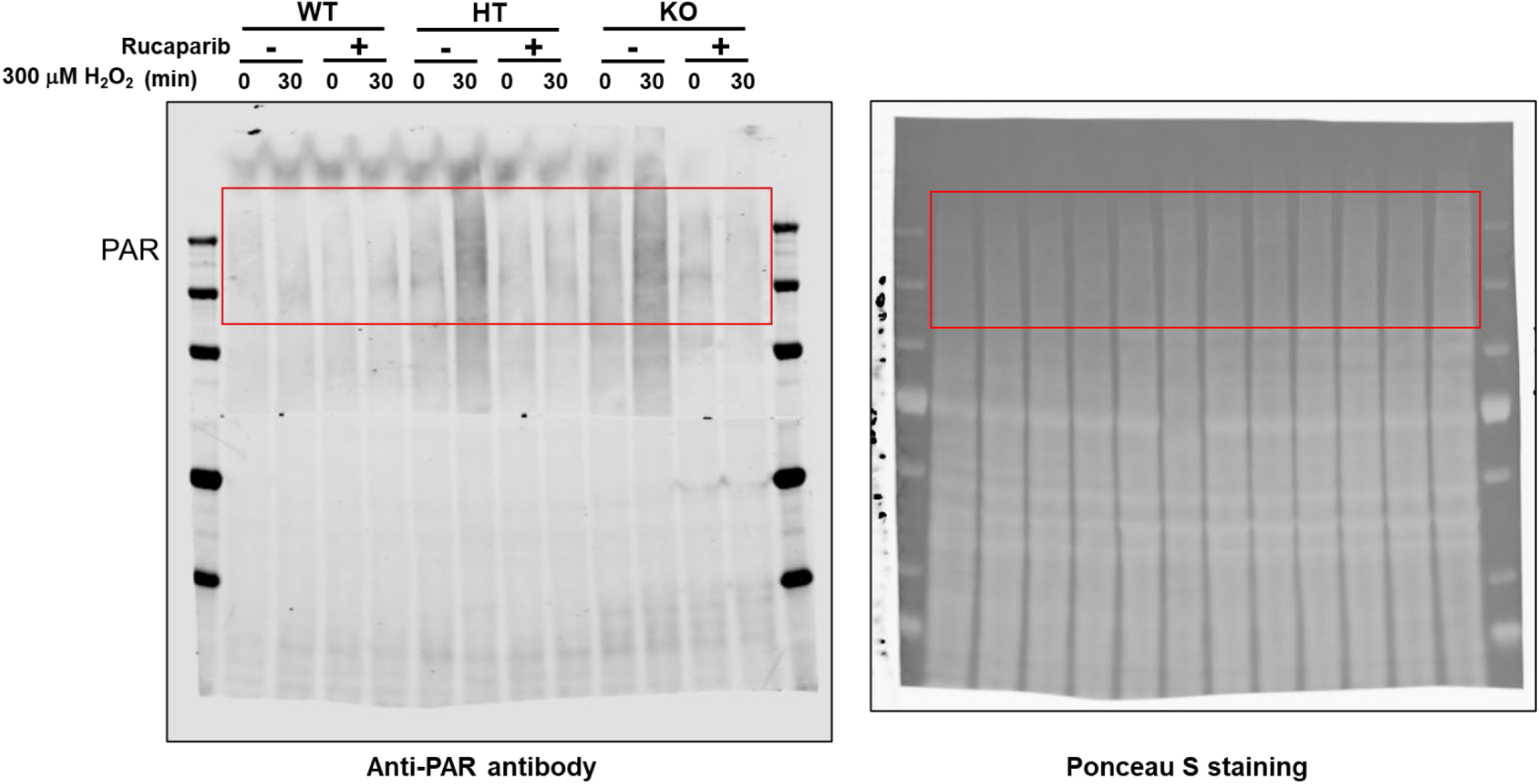

Full unedited gel images for Figure 5E

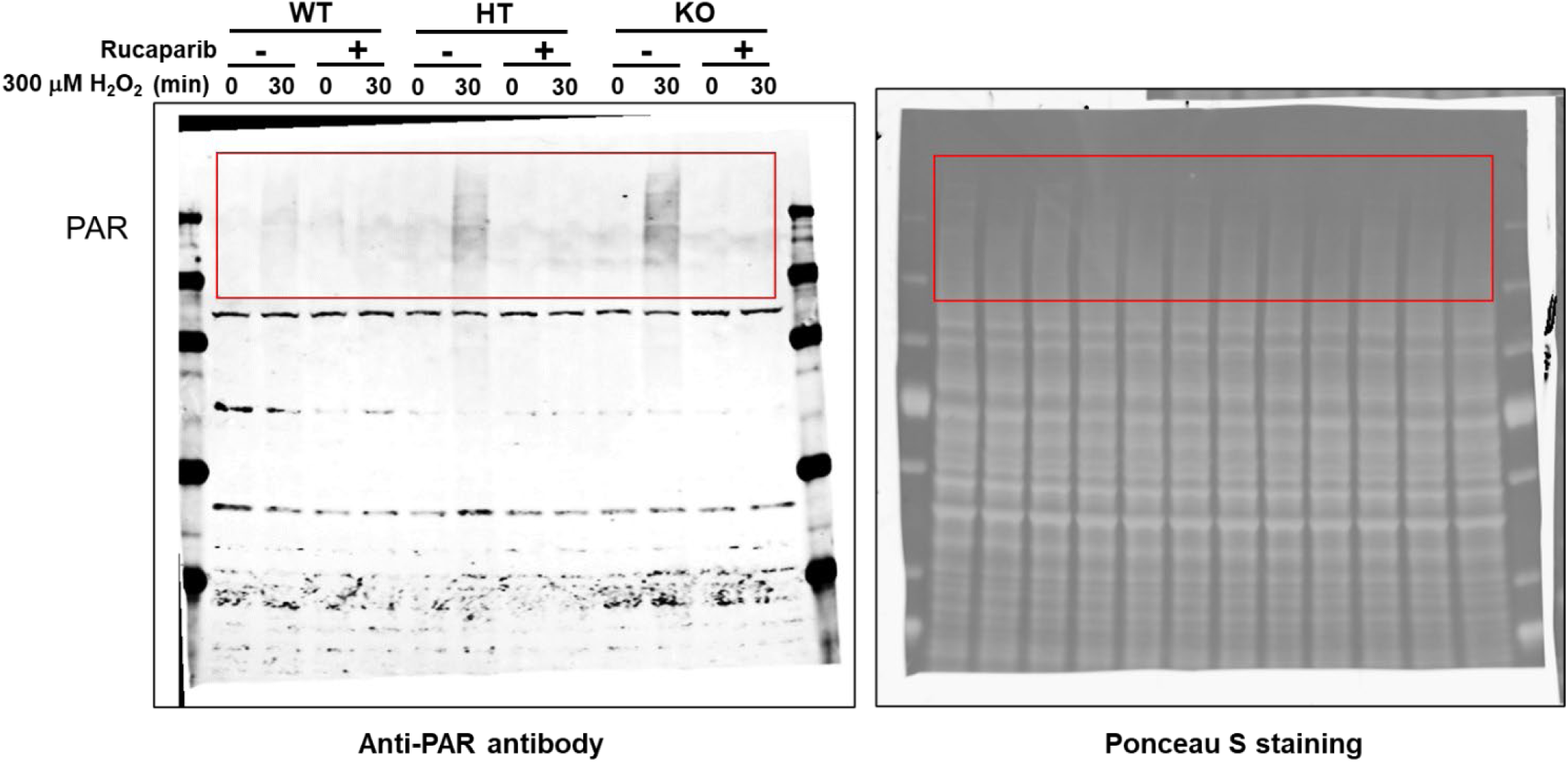

Full unedited gel images for Supplemental Figure 4

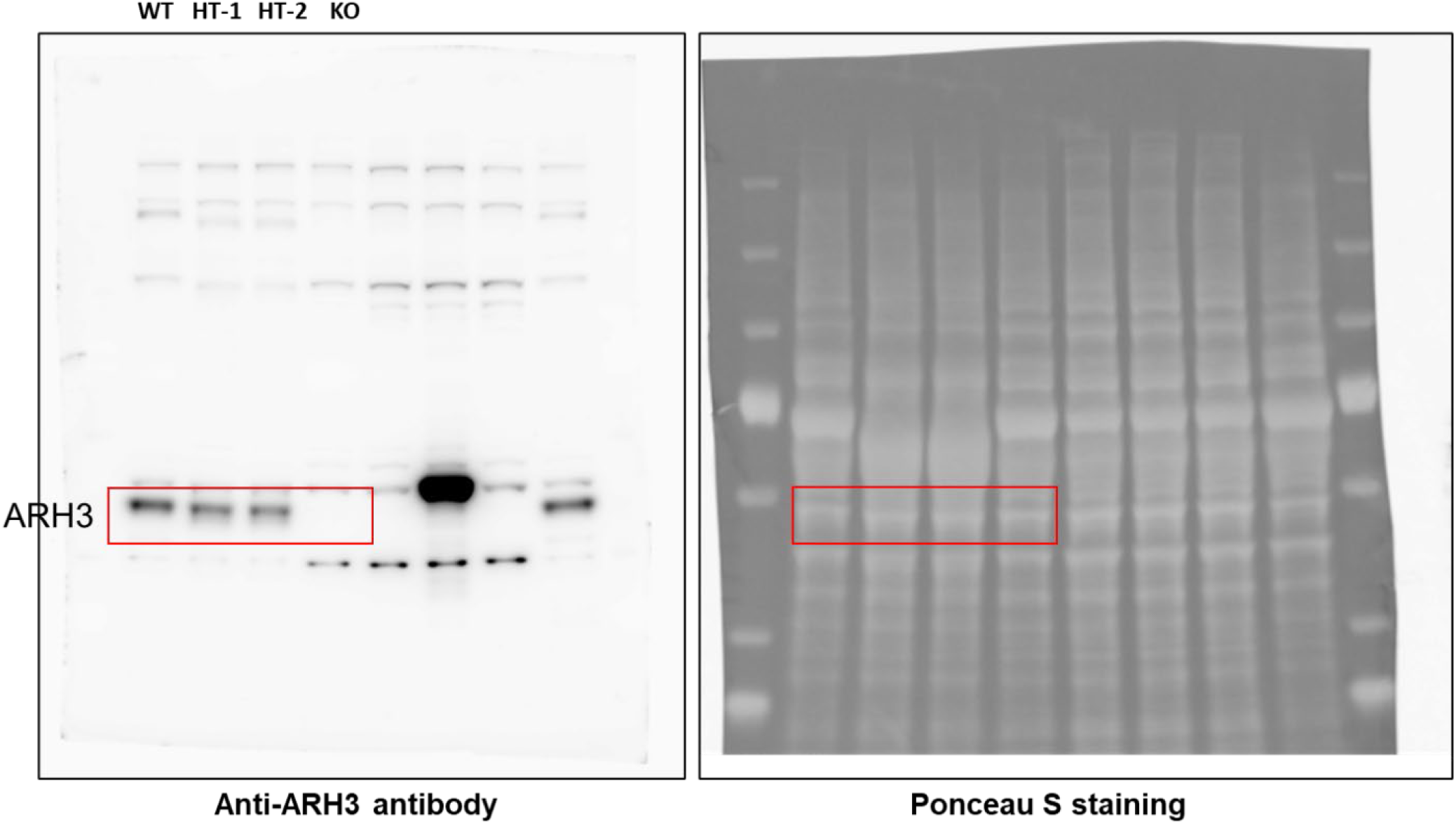

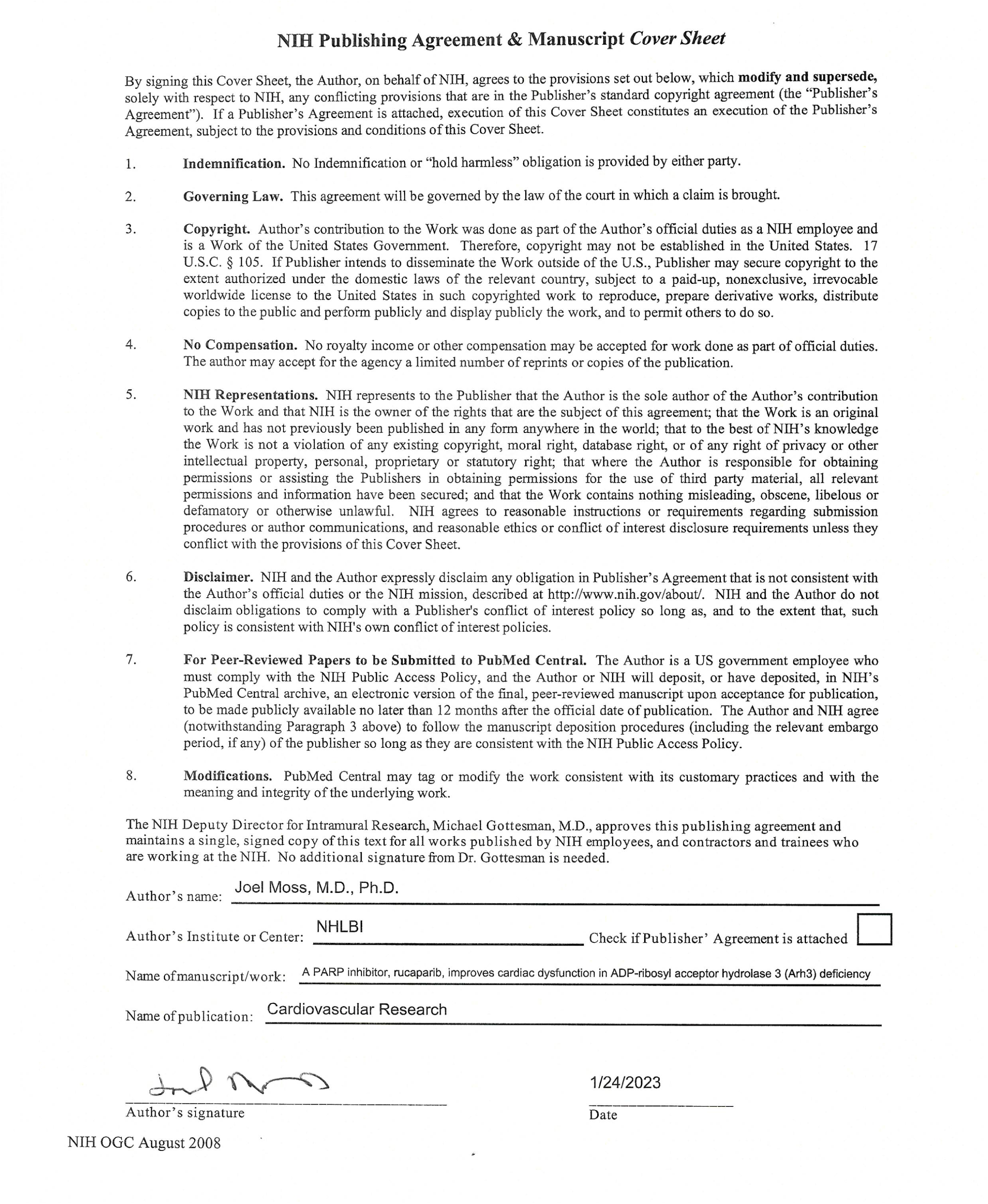

Supplementary video S1

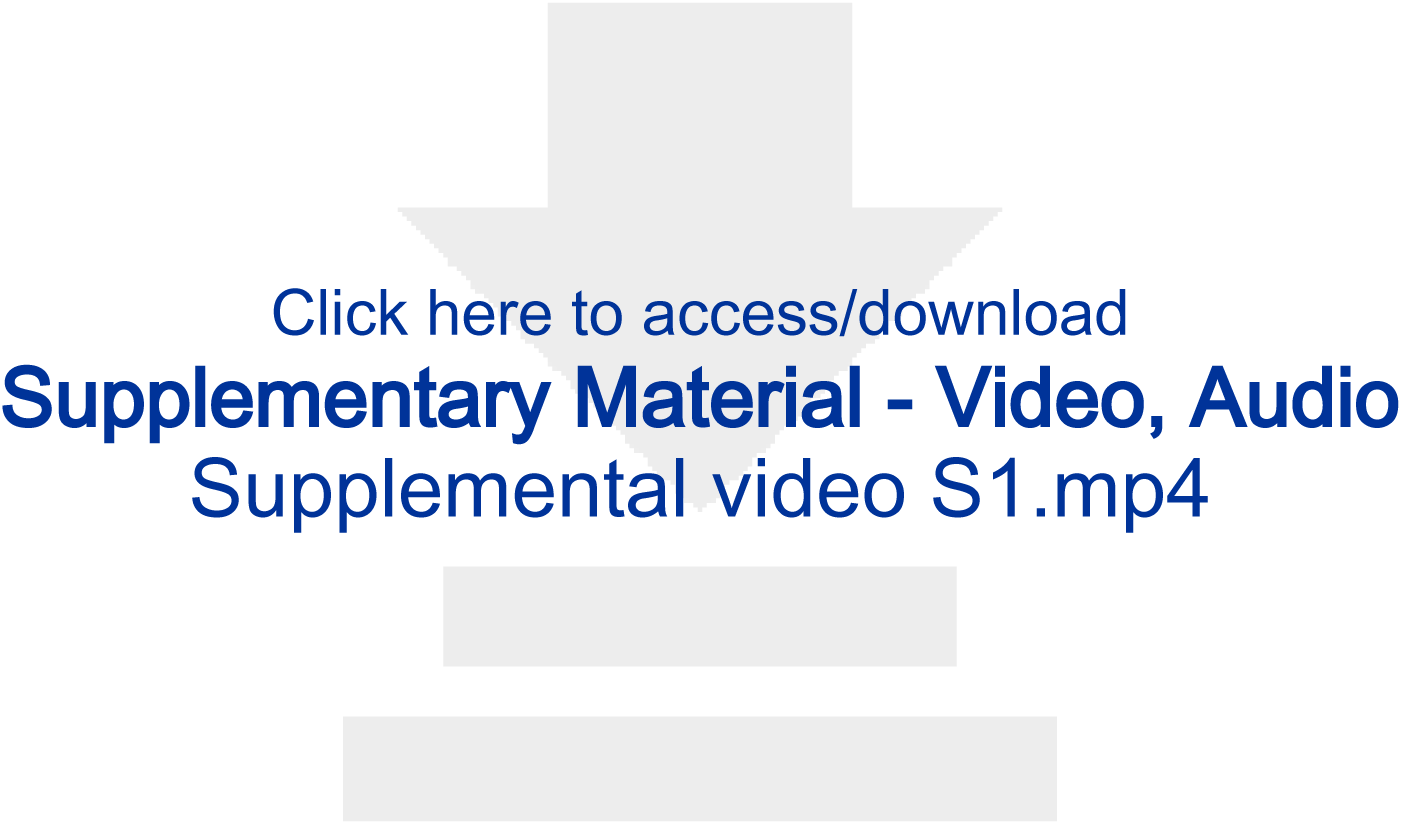

## Notes

### Competing Interest Statement

The authors have declared no competing interest.

